# H3 acetylation selectively promotes basal progenitor proliferation and neocortex expansion by activating TRNP1 expression

**DOI:** 10.1101/2021.03.06.434209

**Authors:** Cemil Kerimoglu, Linh Pham, Anton B. Tonchev, M. Sadman Sakib, Yuanbin Xie, Godwin Sokpor, Pauline Antonie Ulmke, Lalit Kaurani, Eman Abbas, Huong Nguyen, Joachim Rosenbusch, Alexandra Michurina, Vincenzo Capece, Meglena Angelova, Miriam Esgleas, Mareike Albert, Radoslav Minkov, Emil Kovachev, Ulrike Teichmann, Rho H. Seong, Wieland Huttner, Magdalena Götz, Huu Phuc Nguyen, Anastassia Stoykova, Jochen F. Staiger, Andre Fischer, Tran Tuoc

**Affiliations:** University Medical Center, Georg-August-University Goettingen, 37075 Goettingen, Germany; German Center for Neurodegenerative Diseases, 37077 Goettingen, Germany; Max-Planck-Institute for Biophysical Chemistry, 37077 Goettingen, Germany; Ruhr University of Bochum, 44791 Bochum, Germany; Departments of Anatomy and Cell Biology and Stem Cell Biology, Research Institute, Medical University-Varna, Varna 9002, Bulgaria; Institute of Stem Cell Research, Helmholtz Center Munich, German Research Center for Environmental Health, Munich, Germany; Max Planck Institute of Molecular Cell Biology and Genetics, Dresden, Germany; Specialized Hospital for Obstetrics and Gynecology “Prof. Dimitar Stamatov” –Varna, Medical University – Varna, Varna 9002, Bulgaria; Research Center for Functional Cellulomics, Seoul National University, Seoul, Korea; DFG Center for Nanoscale Microscopy & Molecular Physiology of the Brain (CNMPB), 37075 Goettingen, Germany; Zoology Department, Faculty of Science, Alexandria University, Alexandria, Egypt

**Author notes:** Contributed equally (Cemil Kerimoglu, Linh Pham, Anton B. Tonchev, M. Sadman Sakib). Corresponding author: Andre Fischer, Tran Tuoc.

## Abstract

Increase in the size of human neocortex, acquired in evolution, accounts for the unique cognitive capacity of humans. This expansion appears to reflect the evolutionarily-enhanced proliferative ability of basal progenitors (BPs) in mammalian cortex, which may have been acquired through epigenetic alterations in BPs. However, whether or how the epigenome in BPs differs across species is not known. Here, we report that histone H3 acetylation is a key epigenetic regulation in BP amplification and cortical expansion. Through epigenetic profiling of sorted BPs, we show that H3K9 acetylation is low in murine BPs and high in human BPs. Elevated H3K9ac preferentially increases BP proliferation, increasing the size and folding of the normally smooth mouse neocortex. Mechanistically, H3K9ac drives BP amplification by increasing expression of the evolutionarily regulated gene, TRNP1, in the developing cortex. Our findings demonstrate a previously unknown mechanism that controls cortical architecture.

**One Sentence Summary:** H3K9ac promotes basal progenitor amplification, neocortex expansion and gyrification by activating TRNP1 expression in evolution.

## INTRODUCTION

The neocortex of the mammalian brain is radially structured into six neuronal layers and multiple functional domains that form the structural basis for human sensorimotor processing and intellectual ability. During embryogenesis, most cortical neurons are generated from the successive division of neural progenitor cells (NPCs) found in the forebrain germinal zones (i.e., the ventricular (VZ) and subventricular (SVZ) zones). The various types of NPCs can be distinguished by their cell morphology, polarity, ability to generate a given cell lineage, and the site at which they undergo mitosis (*1–6*). The two main types of NPCs in the developing cortex are the apical progenitors (APs) and basal progenitors (BPs). APs include the apical/ventricular radial glia cells (a/vRGs), which divide at the surface of the apical VZ. BPs, which are derived from APs, include the basal (or outer) radial glia (bRGs) and the basal intermediate progenitors (bIPs); the latter lack apical contact and have defined mitotic activities in the inner and outer subventricular zones (iSVZ and oSVZ, respectively) (*1–6*). The aRGs and bRGs are capable of asymmetric division to self-renew, and can directly or indirectly (via BPs) produce neurons (*1–5, 7*).

In most lissencephalic species like mice, most BPs are neurogenic and transient transit-amplifying bIPs (*7, 8*). In many gyrencephalic species, such as human, BPs (including bIPs and bRGs) are capable of undergoing self-amplification through symmetric proliferative divisions before they terminally divide to generate neurons (*1–5, 9*). The intricate folding (gyrification) of the human neocortex is considered to be an evolutionary adaptation to the massive expansion of neuronal populations arising from the high proliferative competence of human NPCs, especially BPs (*2, 3, 10, 11*).

Recent cell sorting- and single cell-based transcriptional profiling analyses have identified a number of factors that are important for BP proliferation, cortical expansion, and folding (*10–21*). Epigenomic methods have been recently established to unravel epigenetic landscapes at the single-cell level, but such strategies are limited by their low coverage of the genome and tend to cluster cells in a manner that is biased toward easily profiled genomic regions. Thus, epigenetic profiling of BPs is still challenging, and the epigenetic mechanisms that are thought to coordinate the expression/repression of gene sets during neocortex expansion in evolution remain unexplored.

Here, we used cell sorting and a new mass spectrometry-based epigenetic profiling to identify H3K9 acetylation as a key epigenetic regulation in BP proliferation, cortical expansion, and cortical folding. We found that although the levels of H3K9ac are comparable in murine and human APs, species-specific differences exist in the histone H3 acetylation of BPs, which is low in mouse BPs and high in human BPs. Interestingly, elevation of H3K9ac in the developing mouse cortex led to BP-specific increases in the promoter H3K9 acetylation and expression of TRNP1, which is a well-known regulator of NPC proliferation and cortical expansion (*11, 22*). The experimental enhancement of H3K9ac also dramatically augmented the TRNP1 expression-dependent proliferative capacity of BPs, leading to enlarged cortical size and folding of the developing mouse cortex. The use of an epigenome editing-based approach to increase H3K9ac specifically at the TRNP1 promoter resulted in increased TRNP1 expression and BP proliferation. Notably, the promoter H3K9 acetylation and expression of TRNP1 were both higher in human BPs compared to mouse BPs, underscoring the relevance of H3K9ac-associated TRNP1 expression changes in neocortical evolution. Our findings demonstrate a novel mechanism of cortical expansion during evolution and suggest that it may contribute to the formation of neocortical gyri in higher primate/human brain.

## RESULTS

### Assessing the epigenetic changes in BPs during cortical evolution

To test whether cortical expansion in evolution is correlated with alteration of the epigenetic landscape, we first investigated whether histone post-translational modifications (PTMs) differ between TBR2 positive (+) BPs from mouse and human cortices (Fig. 1A). To purify TBR2+ BPs from mouse and human developing cortices, we adapted a previously reported intracellular immunofluorescent staining and FACS protocol (*23*). We used an antibody to label intracellular TBR2 in single-cell suspensions isolated from E13.5 and E16.5 mouse cortices and gestational week (GW)14 and 18 human cortices, and then performed cell sorting of TBR2+ and TBR2 negative (−) cells (Fig. 1B/C, fig. S1). Previous studies showed that TBR2 is expressed in bIPs (*24*) and in a subset of PAX6+, HOPX+ bRGs (*25, 26*) in the lissencephalic rodent brain. In the gyrencephalic brains of ferret and primates, TBR2 labeling was seen for bIPs and almost half of the SOX2+, PAX6+ bRG population (*27, 28*). The expression of TBR2 however was not found in TNC+, PTPRZ1+ bRG subpopulation (*14*). Thus, the sorted TBR2+ cells from mouse and human developing cortices actually represent the majority of mouse BPs and human BPs, with the latter including human bIPs and a subset of human bRGs.

**Figure 1.**
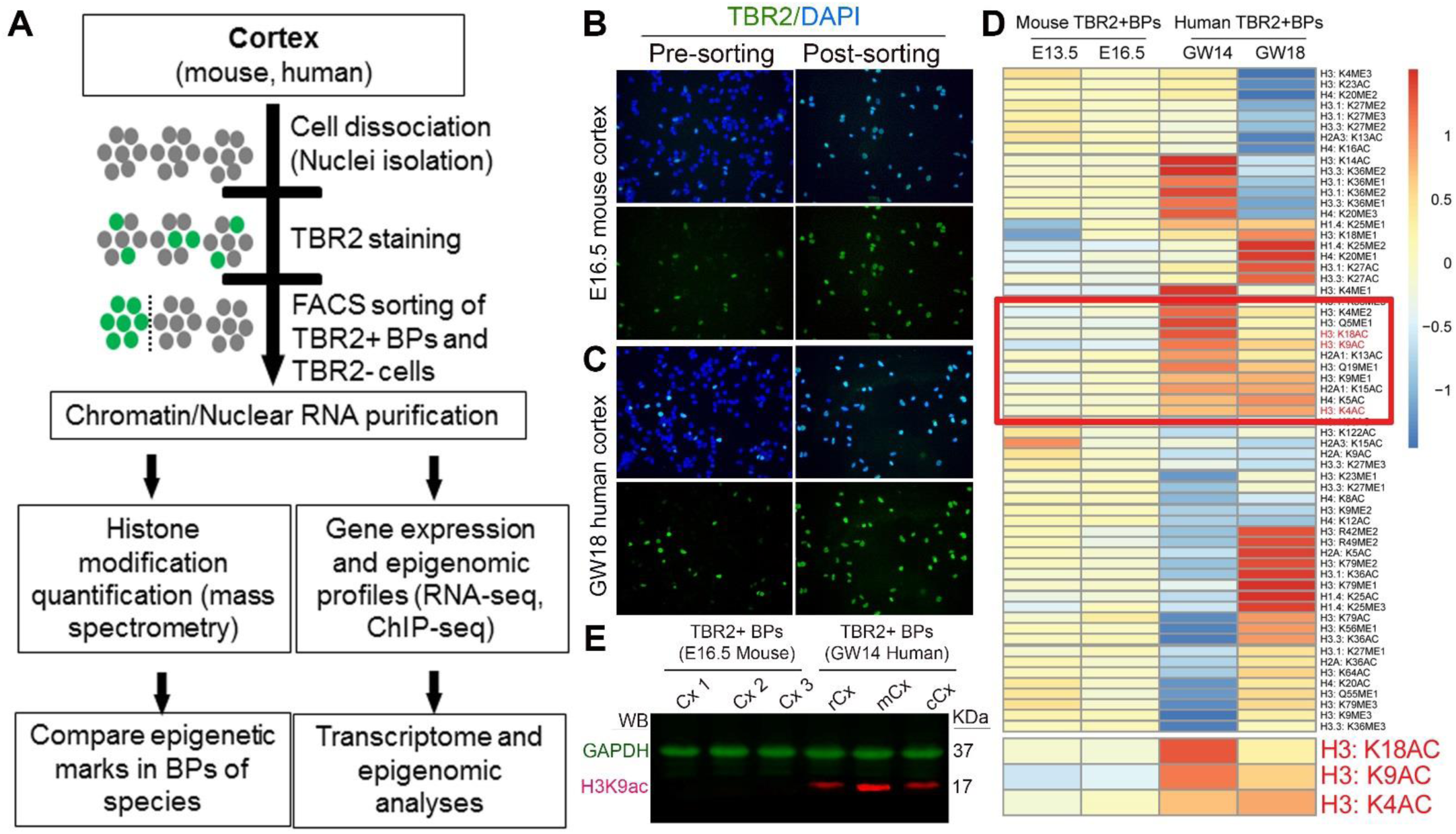
Systematic screening for epigenetic marks showing differential levels between mouse and human basal progenitors (BPs) (A) A scheme of the experimental design used to compare the transcriptomes and epigenomes of mouse and human BPs. (B, C) Purification of TBR2+ BPs in developing mouse and human cortex. Representative images of cell suspensions from mouse cortex (B) and human cortex (C) stained with DAPI and a TBR2 antibody. (D) The data for epigenetic marks is presented as a heat map. Bottom: enlarged pattern showing that the levels of H3K18ac, H3K9ac, and H3K4ac are low in mouse BPs and high in human BPs. (E) Western blot analysis of protein extracts from TBR2+ BPs of whole mouse cortex (Cx) and from human cortex at rostral (r), medial (m) and caudal (c) areas at indicated stages with H3K9ac antibody (in red) and with GAPDH antibody (in green, as loading control).

A newly-established mass spectrometry-based method (see Materials and methods) was applied to quantify peptides containing methylated (me) or acetylated (ac) amino acid residues (Lysine, K; Glutamine, Q; Arginine, R) on the core histones (H2, H3, H4) and the linker histone H1 (Fig. 1D). Intriguingly, the levels of several epigenetic marks, predominantly modified H3 and its variants, appeared to be higher in human BPs at GW14 and GW18 than in mouse BPs at E13.5 and E16.5 (Fig. 1D, in selected frame). Some of the relevant H3 acetylation marks (Fig. 1D; red-labeled in the frame with a higher magnification shown at the bottom), including H3K9ac, H3K18ac, and H3K4ac, were previously shown to be enriched at promoters and enhancers, and to activate transcription (*29, 30*). These findings suggest that epigenetic landscapes, particularly H3 acetylation, reflect extensive changes in BPs during cortical evolution, from rodents to humans.

### Differential levels of acetylated histone H3 in basal progenitors of developing mouse and human cortex

Among the histone acetylation marks found to differ between murine and human BPs, we selected H3K9ac for further in-depth analysis because it exhibited the highest difference between BPs from these species (Fig. 1D) and has been implicated in neuronal function, development, and plasticity (*29, 30*). To validate the screening result, we first performed immunohistochemical (IHC) analyses with antibodies against H3K9ac and progenitor subtype markers (as indicated in the legend of Fig. 2). In E15.5 and E16.5 mouse cortices, high levels of H3K9ac (H3K9ac^high^) were associated with cells in the cortical plate (CP) and the majority of PAX6+ APs in the VZ, whereas fairly low levels of H3K9ac (H3K9ac^low^) were identified in most cells of the intermediate zone (IZ) and many cells of the VZ/SVZ (Fig. 2A/C). Remarkably, all TBR2+ BPs and 58.33±17.48% PAX6+ BPs in the SVZ and IZ were H3K9ac^low^ cells (Fig. 2A/C/D).

**Figure 2.**
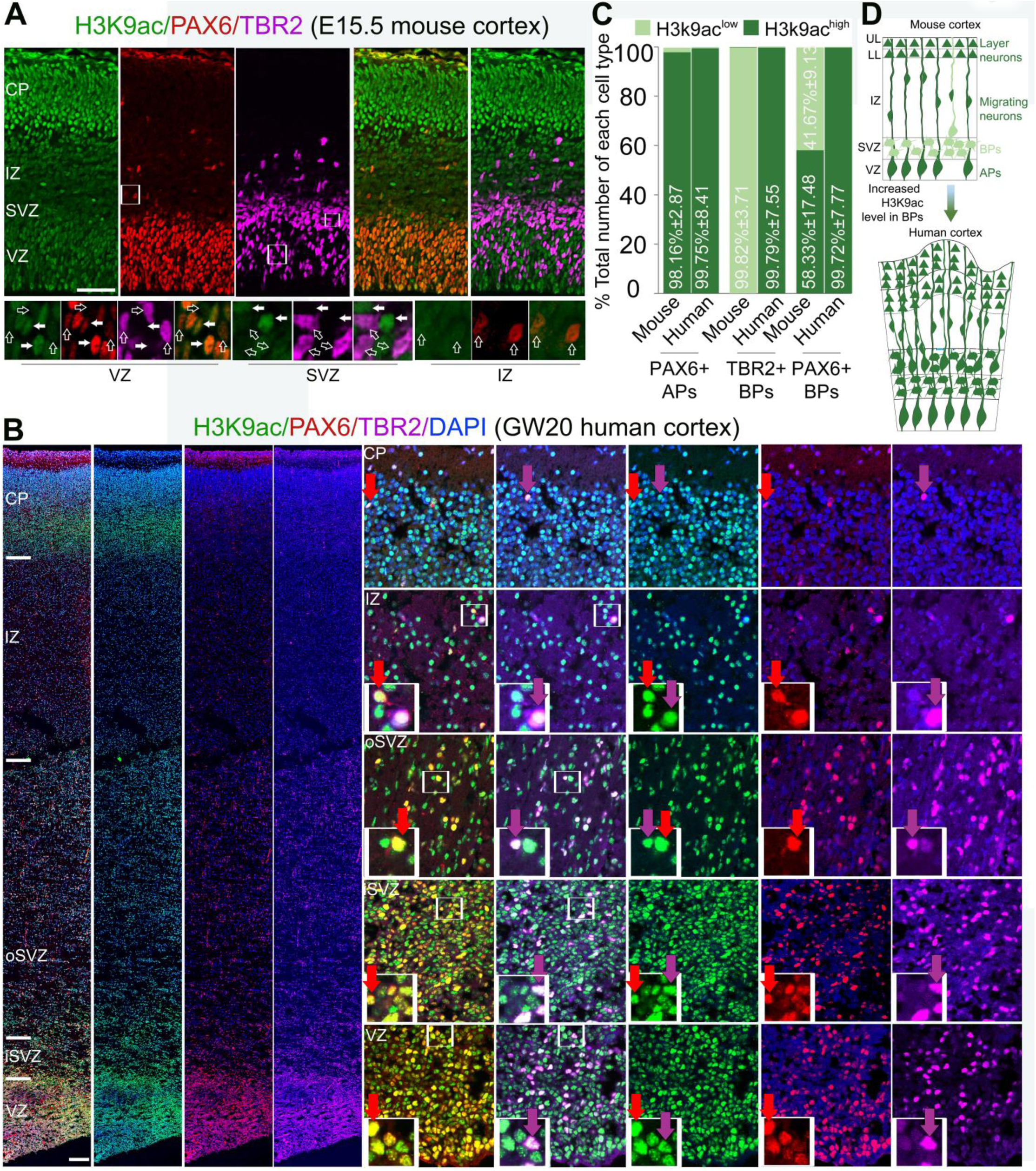
Histone H3K9 is acetylated differently in BPs in the murine and human developing cortex. (A) Images of cortical sections that were obtained from E15.5 mouse embryos and subjected to triple IHC of using antibodies against H3K9ac and PAX6 (to label APs in the VZ and BPs in the IZ) and TBR2 (to mark BPs in the SVZ). The lower panels show higher-magnification images of the boxed areas within the VZ and IZ; they reveal that PAX6^high+^, TBR2^low+^ APs are H3K9ac^high+^ cells (filled arrows), whereas PAX6^high+^ BPs in the IZ and TBR2^high+^ BPs are H3K9ac^low+^ cells (hollow arrows). (B) Images of cortical tissue from human embryos obtained at GW20 and subjected to triple immunolabeling with antibodies against H3K9ac, PAX6, and TBR2. Most of the PAX6+ APs and BPs (red arrows) and the TBR2+ BPs (magenta arrows) are highly immunoreactive for H3K9ac. (C) Statistical analyses of IHC results (shown in A and B) comparing the levels of H3k9ac (H3k9ac^high^ and H3k9ac^low^) in progenitor subtypes in developing mouse and human cortex. The following were compared: PAX6+ APs in the VZ, TBR2+ BPs in the VZ/SVZ, and PAX6+ BPs in the IZ of the mouse cortex; and PAX6+ APs in the VZ, TBR2+ BPs in the VZ/iSVZ/oSVZ, and PAX6+ BPs in the iSVZ/oSVZ of the human cortex. (D) Schema illustrating the higher level of H3 acetylation found in human versus mouse BPs. Abbreviations: APs, apical progenitors; BPs, basal progenitors; VZ, ventricular zone; SVZ, subventricular zone; iSVZ, inner subventricular zone; oSVZ, outer subventricular zone; IZ, intermediate zone; CP, cortical plate. Scale bars = 50 μm.

Likewise, in the human cortex at GW20, a high level of H3K9ac was detected in the CP and in PAX6+ APs (Fig. 2B–D). In striking contrast to the mouse cortex, most PAX6+ BPs and TBR2+ BPs in the human iSVZ and oSVZ were H3K9ac^high^ cells (Fig. 2B–D). The difference in the H3K9ac levels of mouse and human TBR2+ BPs was also confirmed by Western blot analysis with lysates from FACS-collected TBR2+ cells (Fig. 1E).

To determine whether the detected difference in the H3 acetylation levels of the two species was restricted to lysine (K) 9 or also present on other amino acid residues, we examined the pan-acetylation of H3 (H3ac) (fig. S2). Triple IHC analysis of cortical tissues from E15.5 mouse and GW20 human embryos showed that in the mouse cortex, H3ac is highly expressed in PAX6+ APs, but not in TBR2+ BPs. In the human cortex, however, PAX6+ APs in the VZ and PAX62+/TBR2+ BPs in the iSVZ and oSVZ showed high-level H3ac expression (fig. S2A‒C). Notably, numerous KI67+ cycling progenitors were H3ac^low/negative^ and H3K9ac^low/negative^ cells, suggesting that the elevation of H3ac and H3K9ac is not common to all progenitors in human cortex (fig. S2D/E).

Interestingly, our gene ontology (GO) analysis for RNA-seq data of TBR2+ BPs vs. TBR2-cells revealed high expression of many genes encoding for proteins with deacetylase activity, especially histone deacetylases (such as HDAC1, HDAC2, HDAC3, HDAC8, HDAC9, fig. S2F-H). This observation suggests that the high expression of HDACs might remove H3K9ac and possibly other H3ac(s) in TBR2+ BPs in developing mouse cortex.

As histone H3 is acetylated more highly in human BPs than in mouse BPs, we designed experiments to elucidate the *in vivo* outcome of increased H3ac levels, using HDAC inhibition and overexpression of the H3K9 acetyltransferase, KAT2A/GCN5, during mouse cortical development.

### Increased H3 acetylation promotes the generation of basal progenitors

We first tested whether increased acetylation of H3 could increase the genesis and proliferation of BPs in mice. To elevate H3ac, we administered Trichostatin A (TSA), which is a selective class I/II histone deacetylase inhibitor (HDACi), to embryos from control (WT) mice (Fig. 3A). We previously showed that the cortex-specific loss of BAF155, as seen in BAF155cKO embryos, promoted delamination of APs, increasing the population of PAX6+, TBR2+ BPs in the IZ, and diminished the pool of PAX6+ APs in the VZ (*31*) (see also Fig. 3B/C, panels: Control + Veh and BAF155cKO + Veh). As PAX6+ BPs are relatively rare in the WT mouse cortex (*25, 26*), we used the BAF155cKO mutant as a mouse model to investigate the effect of HDAC inhibition on the proliferation of BPs. TSA was injected daily beginning at 12.5 days post coitum (d.p.c.), and WT and BAF155cKO embryos were examined at E16.5–E18.5 (Fig. 3A). TSA treatment had no major effect on the pool of PAX6+/AP2γ+/SOX2+ APs in the VZs of WT and BAF155cKO mutants when compared to the corresponding vehicle (Veh)-treated controls (Fig. 3B/C). Interestingly, TSA injection increased the numbers of PAX6+/AP2γ+/SOX2+/TBR2+ BPs in the SVZ/IZ (Fig. 3B/C, left panels) of WT mice and more pronouncedly in BAF155cKO embryos (Fig. 3B/C, right panels).

**Figure 3.**
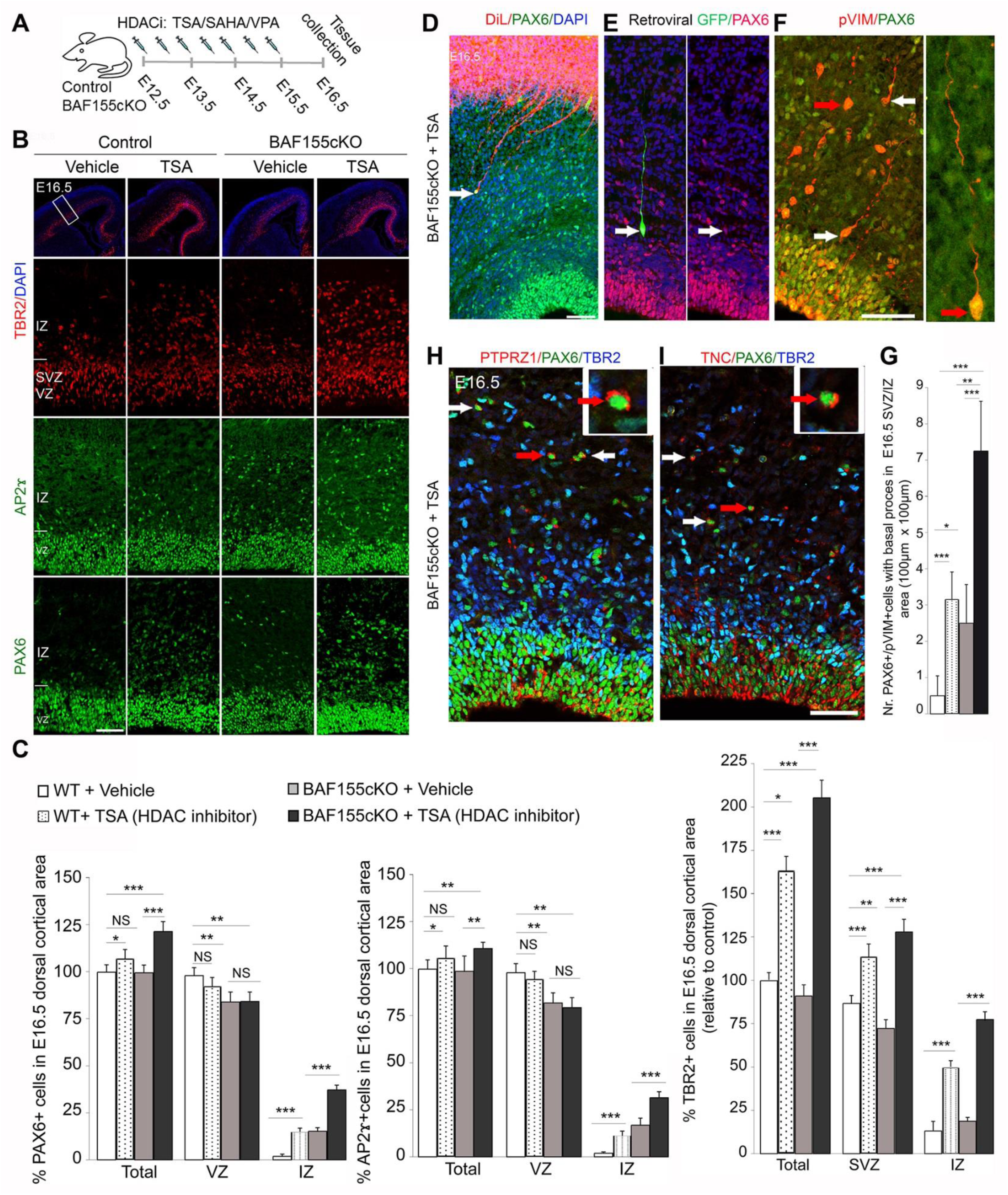
Enhanced acetylation of H3 increases the number of basal progenitors. (A) Experimental paradigm in which BAF155cKO and control embryos were treated with HDAC inhibitors (HDACi: TSA, SAHA, VPA) at the indicated developmental stages. (B) IHC images showing TBR2 (BPs), PAX6 (APs, BPs), and AP2**<u>γ</u>** (APs, BPs) in sections from control and BAF155cKO embryos with or without TSA treatment. (C) Statistical comparisons indicate that increased H3 acetylation enhanced the number of BPs (TBR2+, PAX6+, AP2**<u>γ</u>**+ cells of the IZ) in TSA-treated WT and BAF155cKO embryos compared to vehicle (Veh)-treated controls. (D‒F) Confirmation of BPs/bRGs (arrows) based on their morphology. BPs/bRGs exhibit long basal processes but no apical processes in TSA-treated cortex, as revealed by double labeling of Pax6 with either DiI (D), or retroviral GFP (E), or pVIM (F). Notably, fanned fibers of PAX6+ BPs/bRGs were observed when DiI labeling was applied at the pia (D). (F) High-magnification image of a PAX6+/pVIM+ BPs/bRG, indicated by a red arrow. (G) Statistical analysis comparing number of PAX6+/pVIM+ BPs/bRGs with basal processes in the SVZ/IZ of the indicated embryos. (H, I) IHC analysis shows the expression of the human-enriched bRG markers, PTPRZ1 (H), and TNC (I), in PAX6+/TBR2- cells (arrows) of TSA-treated BAF155cKO cortex at E16.5 (see also fig. S8 for E18.5 cortex). Values are presented as means ± SEMs (\**p* < 0.05, \*\**p* < 0.01, \*\*\**p* < 0.005). Abbreviations: VZ, ventricular zone; SVZ, subventricular zone; IZ, intermediate zone. Scale bars = 50 μm.

Triple IHC analysis of PAX6, TBR2, and KI67 at E16.5 and E18.5 revealed that inhibition of HDAC leads to regionally restricted increases in TBR2+, PAX6+, KI67+ BPs in sections taken from the rostral, middle, and caudal dorsolateral cortex (d/lCx), but not the medial cortex (mCx) (fig. S3). HDAC inhibition exerted a dose-dependent effect, as more TBR2+/PAX6+ BPs were found in E18.5 cortex treated with TSA for 6 days (E12.5–E17.5) compared to those of embryos treated for only 3 days (E12.5–E15.5) (fig. S4A/B). Most TBR2+ BPs of the WT cortex were previously reported to be negative for PAX6 immunostaining (*24*). Interestingly, the proportion of cortical progenitors expressing both PAX6 and TBR2 was high in TSA-treated cortex (fig. S4C), as also reported in the developing gyrencephalic neocortex (*27, 28*).

Abundant PAX6+/DiI+ or PAX6+/retroviral GFP+ or PAX6+/pVIM+ cells with basal processes characteristic of BPs/bRGs were found in TSA-treated mouse cortex (Fig. 3D-G). Notably, TSA treatment resulted in increased numbers of PAX6+/pVIM+ BPs/bRGs, with the highest numbers found in TSA-treated BAF155cKO cortex (Fig. 3G). In addition, cell-surface proteins enriched in human bRG cells, such as TNC and PTPRZ1 (*14*), were also highly expressed in a subset of PAX6+/TBR2- a/bRG cells from TSA-treated mouse cortex (Fig. 3H/I, see also fig. S8).

To further confirm that the genesis of BPs is increased upon HDAC inhibition, pregnant mice were treated with agents whose properties are similar to TSA; these included valproic acid (VPA) and suberoylanilide hydroxamic acid (SAHA), which also inhibit class I/II HDACs. Both VPA- and SAHA-treated developing cortices had more BPs than vehicle-treated cortex (fig. S5; see also Fig. 7H and fig. S10G/H for cortical phenotype of the H3K9 acetyltransferase KAT2A overexpression), supporting the idea that the HDAC inhibition-induced elevation of H3 acetylation directly contributes to increasing the generation of BPs.

### Increased H3 acetylation preferentially enhances the proliferation of basal progenitors but not apical progenitors

The presence of more BPs in TSA-treated BAF155cKO cortex (Fig. 3A-C) suggested that the delaminated progenitors undergo self-amplification in response to enhanced H3 acetylation. To ascertain if H3 acetylation is important for the proliferation of cortical progenitors (APs, BPs), we examined their proliferative capacity by performing IHC with antibodies against PAX6, TBR2, and phosphorylated histone H3 (pHH3) (Fig. 4A/B). Based on the expression of PAX6 in the VZ and pHH3 at the apical VZ surface (Fig. 4A/C), our data suggest that TSA injection did not influence the proliferation of APs, which already have a high endogenous level of acetylated H3. Strikingly, upon HDAC inhibitor treatment, both control and BAF155cKO cortices presented more non-apical proliferating (pHH3+ BPs) cells, along with higher ratios of proliferating BPs/bIPs (TBR2+/pHH3+) to total TBR2+ cells and proliferating BPs/bRGs (basally located PAX6+/pHH3+) to total PAX6+ BPs/bRGs (Fig. 4A–C). Consistent with these data, the numbers of actively cycling BPs (KI67+/TBR2+) and BPs (KI67+/PAX6+) in the SVZ/IZ were considerably increased in TSA-treated BAF155cKO cortex (Fig. 4D/E). In addition to the lengthened cell cycle in neural progenitors from primate than that from rodent (*3*), both proliferative APs and BPs exhibit a substantially longer S phase than neurogenic progenitors (*32*). We therefore focused on examining the effect of TSA on progression from S phase to G2-M phases in APs and BPs. Given that the average lengths at mid-gestation stage of the S phase and G2-M cell cycle phases are about 3.5 hours and 2 hours, respectively (*32*), we then examined whether the TSA treatment affected the progression within S-G2-M phases in APs and BPs. A thymidine analog injection paradigm (4 hours-IdU pulse-labeling) was used to mark cortical progenitors within these phases. We performed double immuno-staining with antibodies against IdU and pHH3 to label APs (apical surface-located IdU+/pHH3+) and BPs (basally located IdU+/pHH3+), which already entered G-M phases (Fig. S6A/B). Remarkably, fewer BPs (but not APs) reached late G2-M phases in TSA-treated cortices compared to Veh-treated cortices. This implicated that TSA treatment resulted in lengthening of cell cycle progression in S-G2-M phases in BPs, specifically.

**Figure 4.**
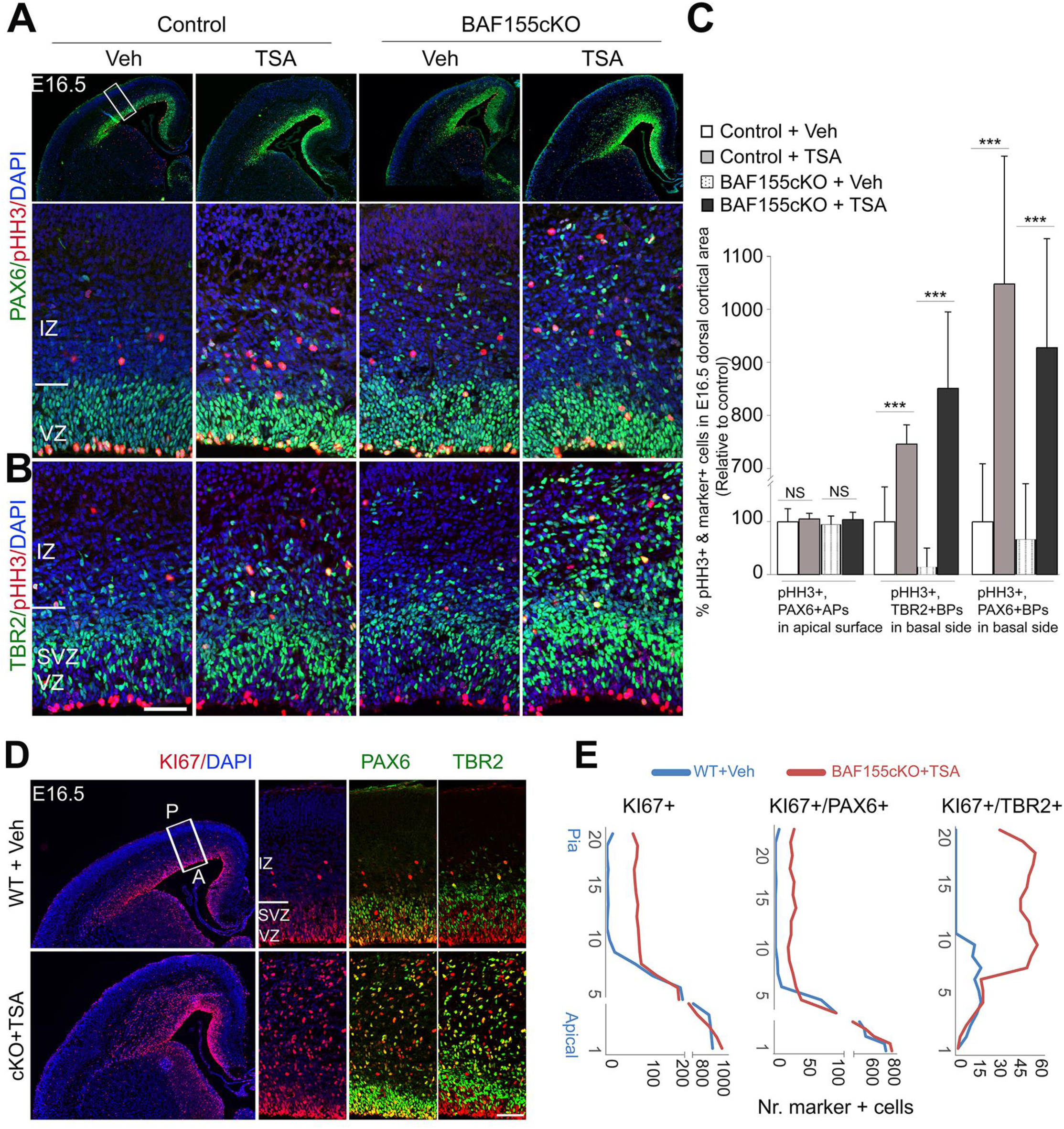
H3 acetylation specifically promotes proliferation of basal progenitors in developing mouse cortex. (A, B) Images showing double IHC of PAX6/pHH3 and TBR2/pHH3 in control or BAF155-deficient cortex treated with Veh- or HDACi, as assessed at E16.5. Lower panels show higher magnifications of the areas indicated by the white box. (C) Statistical analyses reveal that elevated acetylation of H3 causes a significant increase in the number of TBR2+/pHH3+ BPs and PAX6+/pHH3+ BPs, whereas the number of apical PAX6+/pHH3+ APs cells is not affected. (D) IHC images for KI67/PAX6 or KI67/TBR2 staining of Veh-treated WT and TSA-treated BAF155cKO cortex at E16.5. Right panels show higher magnifications of the areas indicated by the white box. (E) Quantified distribution of proliferating KI67+/PAX6+ APs/BPs and KI67+/TBR2+ BPs from the apical surface to the pia in vehicle-treated WT cortex and the corresponding region of the TSA-treated BAF155cKO cortex shown in A. Values are presented as mean ± SEM (\**p* < 0.05, \*\**p* < 0.01, \*\*\**p* < 0.005). Abbreviations: VZ, ventricular zone; SVZ, subventricular zone; IZ, intermediate zone. Scale bars = 50 μm.

Together, these data indicate that the proliferative capability of murine BPs is enhanced by elevated acetylation of H3.

### Identification of H3K9ac target genes in TBR2+ BPs

Given that TSA treatment had a much stronger effect on BP genesis in BAF155cKO cortex, we firstly compared the gene expression and genome-wide H3K9ac profiles of TSA- and vehicle-treated BAF155cKO cortices by RNA-Seq and ChIP-Seq (Fig. 5A/B; Table S1, S2). We found that TSA treatment yielded upregulation of 1961 genes and downregulation of 1799 genes (*p*-value < 0.01 & |fold change| > 1.2) (Fig. 5A, Table S1). Examination of H3K9 acetylation on the TSA-regulated genes revealed that the upregulated genes had clear increases in the level of H3K9ac (fig. S7A), whereas the downregulated genes did not show any significant change in H3K9ac (fig. S7B). This result is in accordance with data showing that loss of H3K9 acetyltransferases can trigger activation of a set of genes via secondary effects (*33*).

**Figure 5.**
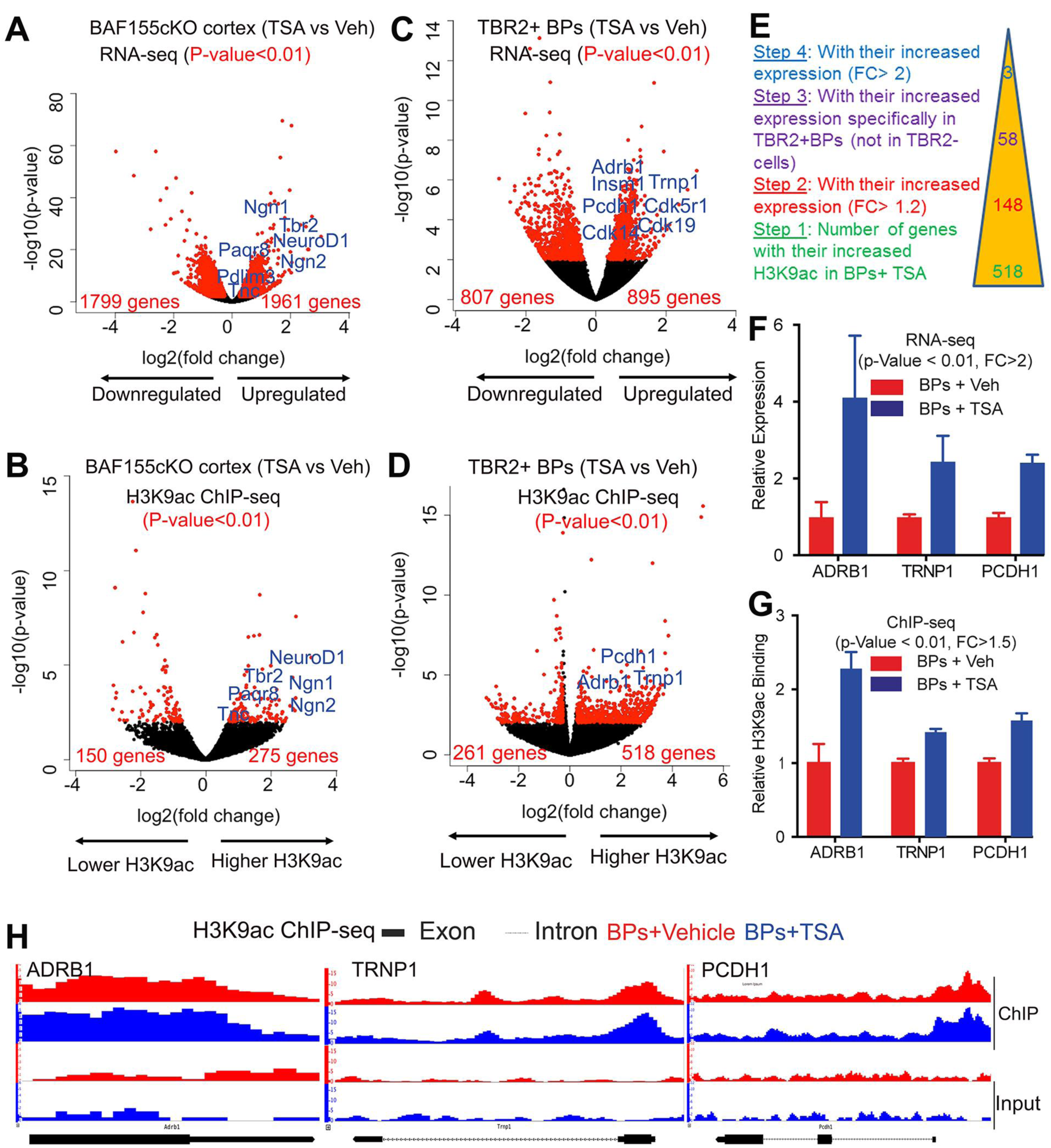
Identification of H3K9ac target genes in TBR2+ BPs. (A–D) Volcano plots showing statistically significant changes (Paired Student’s t-test < 0.01, FC > 1.2) visualized by our RNA-Seq (A, C) and H3K9ac ChIP-Seq (B, D) analyses of E16.5 BAF155cKO cortex (A, B) and TBR2+ BPs (C, D) in TSA vs. Vehicle experiments (see also Fig. S7E/F for TBR2-cells). (E) Stepwise exclusion parameters were applied to the H3K9ac ChIP-Seq and RNA-Seq data sets to search for genes specifically activated by H3K9ac in TBR2+ BPs. (F) The expression levels of ADRB1, TRNP1 and PCDH1 are upregulated in TSA-treated BPs. (G/H) The level of H3K9ac (G) and distribution of H3K9ac along the gene bodies (H) of ADRB1, TRNP1, and PCDH1 in TSA-treated BPs (red) and Vehicle-treated BPs (blue). Input (bottom row) and distributions after immunoprecipitation (upper two rows) are depicted.

In accordance with our IHC data showing an increase in the number of TBR2+/PAX6+ BPs, many neurogenic BP/bIP genes (e.g., NEUROG1, NEUROG2, EOMES/TBR2, NEUROD2, NEUROD6) were upregulated in the cortex of TSA-injected BAF155cKO mutants (Fig. 5A; Table S1). TSA treatment also yielded increased expression of genes previously shown to be enriched in BPs/bRGs (e.g. TNC, PTPRZ1, PAQR8, GIGYF2, PDLIM3, ZC3HAV1; fig. S8A; Table S1) (*14*). The increased expression of many BP-enriched genes in response to HDAC inhibition in both BAF155cKO and WT cortex was also confirmed by qPCR and IHC analyses (fig. S7C, S8). Furthermore, the results from our ChIP-Seq and ChIP/qPCR analyses also confirmed that many of the BP genes (e.g., TNC, PAQR8, NEUROG2, EOMES, NEUROD1, NEUROD6; fig. S7D; Table S2) exhibited increases in their H3K9ac levels in the cortices of TSA-treated BAF155cKO and WT embryos. These findings indicated that H3 acetylation positively regulates the expression of BP genes.

Next, we evaluated the TSA treatment response of TBR2+ BPs (Fig. 5C/D; Table S3, S4) and TBR2-cells (fig. S7E/F; Table S5, S6). For this purpose, TBR2+ and TBR2-nuclei were sorted by FACS (Fig. 1; fig. S1). TSA treatment yielded expressional upregulation of 895 genes in TBR2+ BPs. To provide additional evidence that H3 acetylation induces expression of BP genes during evolution, we compared the upregulated genes in TSA-treated TBR2+ BPs with BP/IP genes that were recently identified to be specific for macaque or human (*34*). Remarkably, TSA treatment provoked expression of 40.1% (166 out of 414) macaque-specific BP genes and 13.3% (62 out of 467) human-specific BP genes in TBR2+ BPs in the developing mouse cortex (fig. S7G/H; Table S7), suggesting that H3 acetylation has an evolutionary relevance in BP genesis during mammalian evolution. Among the upregulated genes in TSA-treated TBR2+ BP genes, 37 showed correlation between their upregulation and an increased level of H3K9ac upon TSA treatment (fig. S7I; Table S4; *p*-value < 0.01). Surprisingly, neither the H3K9ac level nor the expression levels of typical BP genes (e.g., EOMES, NEUROG1, NEUROG2, NEUROG1, NEUROD4) were found to be altered in TSA-treated BPs (fig. S7J; Table S3-S6). Thus, the upregulated expression of most typical BP genes in TSA-treated developing mouse cortex seems to be a secondary effect.

The increased proliferation of BPs, which was visualized as increases in TBR2+/pHH3+ and TBR2+/KI67+ cells (Fig. 4), suggested that the expression of proliferation-related genes would be increased. Indeed, we found that the expression levels of cell cycle genes (e.g., CDK19, CDK5R1, CDK14) and INSM1, which is known to promote BP proliferation, were upregulated in TSA-treated BPs (Fig. 5C; Table S3). Remarkably, however, the levels of H3K9ac at the promoters of these genes were not increased following TSA treatment (Fig. 5D; Table S4). These findings suggested that TSA treatment might influence the expression of factors that act upstream of the aforementioned proliferation-regulated genes.

To identify candidates for functional analysis, we undertook an unbiased approach. From among all genes that exhibited increased H3K9ac at their promoters in TBR2+ BP cells after TSA injection (|fold change| > 1.2; Fig. 5E), we selected those that also showed a trend for increased expression (|fold change| > 1.2; Fig. 5E). This screen netted 148 genes (Fig. 5E). From them, we filtered out those that exhibited increased expression in TBR2-cells, yielding 58 genes that had increased H3K9ac and were uniquely upregulated in TBR2+ BP cells following TSA treatment (Fig. 5E). These genes were further filtered according to their basal expression levels (baseMean > 20) and selection of those that increased by more than 2-fold. This pipeline yielded three candidates: ADRB1, TRNP1, and PCDH1 (Fig. 5F-H, fig. S7K). This suggests that H3 acetylation promotes BP amplification by directly activating the expression of a small set of genes.

### H3 acetylation controls the amplification of basal progenitors by regulating TRNP1 expression in the developing cortex

In the developing mouse cortex, TRNP1 is exclusively localized in the nuclei of APs in the VZ and nascent neurons in the CP, with little to no TRNP1 expression seen among TBR2+ BPs in the SVZ (Fig. 6A) (*11*); it thus exhibits a pattern similar to that of H3K9ac (Fig. 1E, Fig. 2). In line with this, the expression and promoter H3K9ac levels of TRNP1 were lower in TBR2+ BPs than in TBR2-cells of the developing mouse cortex, as revealed by our RNA-Seq and ChIP-Seq experiments (fig. S9A/B). Notably the expression (Fig. 6A/B) and promoter H3K9ac (Fig. 6C, fig. S9C) levels of TRNP1 were strikingly higher in human TBR2+ BPs than in mouse TBR2+ BPs. These data suggest that the H3K9ac level directly controls the expression of TRNP1 (also see the model in Fig. 7I). This proposal was further corroborated by our observation that TRNP1 expression is increased specifically in TBR2+ cells upon TSA treatment (fig. S9D/E).

**Figure 6.**
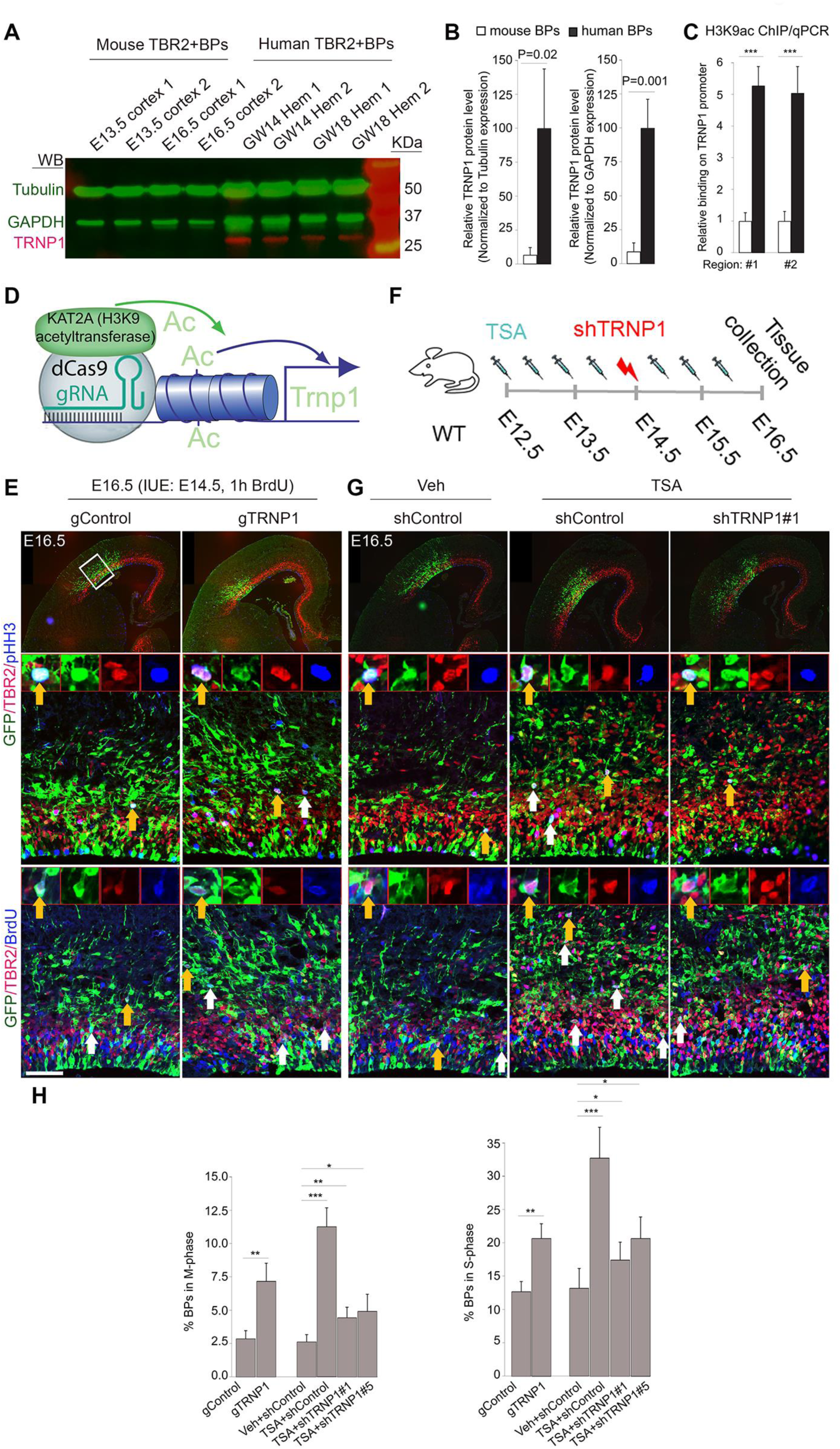
H3 acetylation controls the proliferation of BPs by activating TRNP1 expression. (A, B) Western blot analysis of protein extracts from TBR2+ BPs obtained from mouse and human cortices at the indicated stages, as assessed with antibodies against TRNP1 (red), Tubulin, and GAPDH (green, as loading controls). (B) Relative levels of TRNP1 protein are presented in the diagram. Little to no TRNP1 protein is expressed in mouse BPs, whereas this expression is relatively high in human BPs. (C) ChIP-qPCR comparing the H3K9ac levels at the TRNP1 promoters in mouse and human TBR2+ BPs. (D) Schematic overview of the CRISPR/dCas9-based deposition of H3K9ac at the TRNP1 promoter, which was used to activate its expression. (E) Mouse E14.5 dorsolateral cortex was *in utero* electroporated with a gTRNP1-dCas9-KAT2A-T2A-eGFP plasmid (gTRNP1) or gControl-dCas9-KAT2A-T2A-eGFP plasmid (gControl), and IHC analysis of GFP, TBR2, pHH3, or BrdU was performed at E16.5. Images represent triple optical sections. White box indicates areas shown at higher magnification and red arrows point to examples of cells that are immunoreactive for GFP, TBR2, pHH3, or BrdU. (F) Descriptive scheme of the rescue experiment, in which mouse cortex was treated with the HDAC inhibitor, TSA, and gTRNP1 or shControl constructs at the indicated stages. (G) Triple IHC analysis for the markers listed in panel E. (H) Statistical quantification of the results shown in panels E and G is shown. Values are presented as mean ± SEM (\**p* < 0.05, \*\**p* < 0.01, \*\*\**p* < 0.005). Abbreviations: VZ, ventricular zone; SVZ, subventricular zone; IZ, intermediate zone. Scale bars = 50 μm.

**Figure 7.**
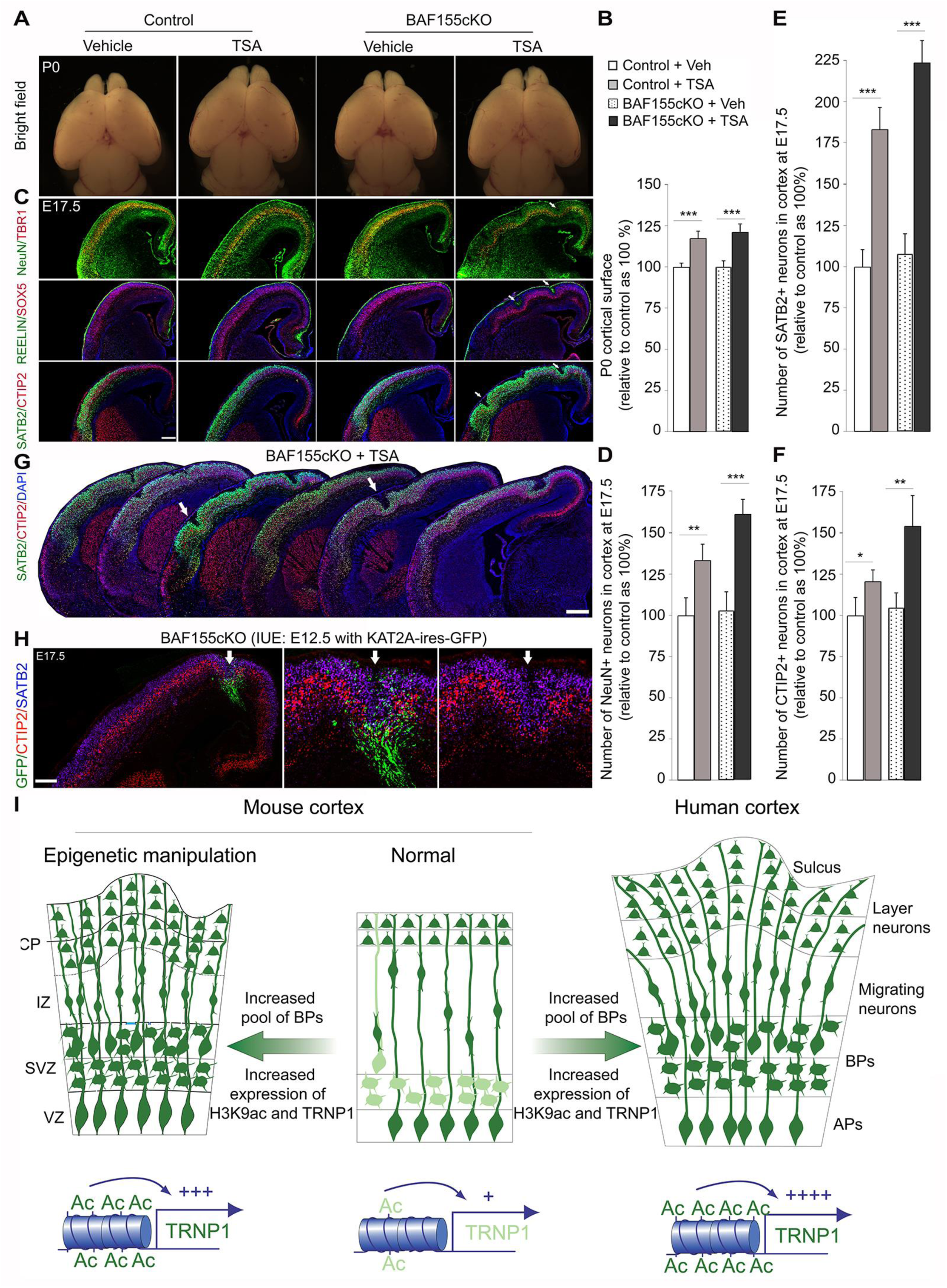
Elevated H3ac levels promote cortical expansion and folding. (A) Dorsal-view images of brains from control and BAF155cKO mice with and without TSA treatment (B) and quantification of their cortical surfaces at P0 (C, see also Supplemental methods). (C-F) IHC (C) and quantitation (D-F) of the number of neuronal subsets labeled by the neuronal markers, SATB2, CTIP2, TBR1, and NeuN, in control and BAF155cKO cortices with and without HDAC inhibitor treatment at E17.5. Elevated H3 acetylation increases neurogenesis and cortical thickness and induces gyrification (indicated by arrows). (G) Serial coronal sections at E17.5 marked by CTIP2 and SATB2 expression showing the folded cortex in different cortical areas. (H) Images of GFP, SATB2, and CTIP2 immunofluorescence of a coronal section of an E17.5 cortical hemisphere from a mouse embryo that had been *in utero* electroporated at E12.5 with KAT2A-ires-eGFP expression plasmids. Lower images show higher-power magnification of a sulcus-like structure, which is indicated by white arrows. (I) Hypothetical model proposing how the changes in H3K9ac levels in evolution and in mouse models with epigenetic manipulation affect the H3K9ac level at the TRNP1 promoter, the expression of TRNP1, the proliferation of BPs, and the expansion and folding of the cortex. Symbols: +, +++, and ++++ indicate weak, moderate, and strong relative expression levels, respectively (Photo Credit: Dr. Tran Tuoc, Ruhr University of Bochum). Abbreviations: VZ, ventricular zone; SVZ, subventricular zone; IZ, intermediate zone; CP, cortical plate; APs, apical progenitors; BPs, basal progenitors; Ac, H3K9ac. Values are presented as mean ± SEM (*p < 0.05, **p < 0.01, ***p < 0.005). Scale bars = 100 μm.

To more directly investigate whether a higher level of H3K9ac at the TRNP1 locus might enhance the gene expression and proliferation of BPs in the mouse developing neocortex, we sought to increase H3K9ac at the TRNP1 locus and examine the associated phenotype. To increase deposition of the H3K9ac mark, we adapted a CRISPR/dCas9- based system (*35*) that allows targeted editing of H3K9ac in the developing cortex (Fig. 6D). Given that the levels of H3K9ac at the TRNP1 promoter differed between mouse TBR2+ BPs and human TBR2+ BPs (Fig. 6C) and between mouse TBR2+ BPs and TBR2-cells (fig. S9B), and TSA treatment increased H3K9ac at the TRNP1 promoter specifically in BPs (fig. S9D/E), we selected the TRNP1 promoter for targeted H3K9ac editing. Several guide RNAs (gRNAs) targeting the TRNP1 promoter were designed and tested (fig. S9F/G). To edit histone acetylation at the TRNP1 locus, we generated plasmid constructs (gTRNP1-dCas9-KAT2A-T2A-eGFP) harboring DNA sequences that encoded a gRNA, dCas9 (a nuclease-deficient Cas9) fused with an H3K9ac writer KAT2A, and a fluorescent reporter (GFP) (Fig. 6D). Two gRNA-expressing constructs (gRNA#2 or gRNA#4), designated as gTRNP1, were used to target the TRNP1 locus (fig. S9F/G).

To validate that H3K9ac levels changed upon introduction of gTRNP1, we carried out ChIP-qPCR experiment using FACS-purified transfected (GFP+) Neuro2A cells. H3K9ac levels were observed to have increased significantly at the TRNP1 promoter region upon gTRNP1 transfection, compared to control conditions (fig. S9H). In addition, qPCR analysis of the sorted cells revealed that TRNP1 expression was also increased following transfection with the generated editing constructs (fig. S9I). These results validate the ability of the CRISPR/Cas9-mediated epigenome editing system in depositing H3K9ac at the TRNP1 promoter and increasing the expression of TRNP1 in neural cells.

To achieve epigenome editing-based modulation of H3K9ac levels in the developing cortex, we delivered the constructs into APs by *in utero* electroporation (IUE) at E14.5. At E16.5, we analyzed the progeny of the electroporated APs, including TBR2+ BPs (Fig. 6D/E). By comparing the proportion of TBR2+/pHH3+ (BPs in M-phase of the cell cycle) and TBR2+/BrdU+ (BPs in S-phase) cells among the targeted TBR2+ BPs (revealed by GFP) in control- and editing construct-injected cortices, we examined whether the altered levels of H3K9ac might influence the proliferation of BPs (Fig. 6E/H). Similar to TSA treatment, epigenome editing-based augmentation of the promoter H3K9ac level of TRNP1 increased the percentage of basal mitosis (GFP+/TBR2+/pHH3+ or BrdU+) among targeted BPs (GFP+/TBR2+; Fig. 6E/H).

Because TRNP1 overexpression was shown to promote the proliferation of NSCs in the developing mouse and ferret cortex (*11, 22*), we performed rescue expression to examine whether a specific increase in TRNP1 expression caused the aberrantly enhanced proliferation of BPs upon TSA treatment. To this end, we electroporated TSA-treated cortex with shTRNP1 constructs (*11*) to downregulate TRNP1 expression (Fig. 6F/G). Remarkably, the observed TSA treatment-induced proliferation phenotype of TBR2+ BPs was largely rescued by TRNP1 knockdown in the cortex (Fig. 6G/H). These findings suggest that the elevated level of H3K9ac at the TRNP1 promoter is directly involved in facilitating the proliferation of TBR2+ BPs (also see the model in Fig. 7I).

### Elevated H3ac leads to radial expansion and induces gyrification of the developing mouse cortex

To address the role of H3 acetylation in late corticogenesis *in vivo*, mouse embryos were treated with TSA for a prolonged period. Remarkably, TSA-treated mice exhibited cortices with larger surface areas compared with vehicle (Veh)-injected controls (Fig. 7A/B). To evaluate the effect of the increased level of H3 acetylation on neuron production in the developing cortex, we examined the expression of laminar and neuronal subtype-specific genes at E17.5 (Fig. 7C). Immunostaining for the pan-neuronal marker, NeuN, indicated that there was a considerable increase in cortical radial thickness after TSA treatment (Fig. 7D). The number of SATB2+ callosal and CTIP2+ subcerebral projection neurons was also significantly increased in the cortex of TSA-treated embryos compared with controls (Fig. 7E/F).

Previous evidence suggested that an increased number of bRG cells offers extra scaffolding for the radial migration of neurons and causes divergence (fanning out) of radial glial processes in gyrated cortices (*2, 4, 10, 11*). Indeed, RC2 and NESTIN immunostaining, which reveal the layout of RG processes, showed that in the TSA-treated cerebral cortex radial fibers spread out divergently when traversing the CP (fig. S10A, white dashed lines). Fanned-out fibers of PAX6+ bRGs were observed by labeling with the lipophilic dye, DiI, which was applied at the pia (Fig. 3D). At E17.5, most early-born neurons (L6, L5) had completed their migration. Strikingly, TSA treatment often led to mild folding of the cortex in WT embryos (fig. S10B-D), but more profound folding in BAF155cKO mutants in distinct cortical areas, as indicated by L6 (TBR1) and L5 (CTIP2) immunostaining (Fig. 7G, S10B–D). Although HDAC inhibitor-treated mice died soon after birth, increased folding of TSA-treated cortex was observed at all examined stages (E16.5–P0) (fig. S10D). Our findings further corroborate the concordance between abundance of BPs and folding feature of the mammalian cortex as the inhibition of HDAC leads to regionally restricted increases in BPs (fig. S3) and cortical folding (fig. S10B) in sections taken from the rostral, middle, and caudal dorsolateral cortex (d/lCx), but not the medial cortex (mCx). We distinguished cortical folding by basement membrane intactness (fig. S10E/F) and rostro-caudal continuity of both sulci and gyri; this allowed us to distinguish our findings from the defining features of classical neuronal ectopias, such as that seen in lissencephaly type II (also called cobblestone lissencephaly).

KAT2A and KAT2B are H3K9 acetyltransferases as the dual deletion of KAT2A and KAT2B led to the complete elimination of H3K9ac (*33, 36*). Because the loss of KAT2A in NSC-specific mouse mutants caused microcephaly (*37*) and KAT2A directly involved in H3K9 acetylation at TRNP1 promoter (Fig. 6D/E, Fig. S9G-I), we investigated whether increasing the level of H3K9ac by overexpressing KAT2A might induce proliferation of BPs and cortical folding. E13.5 WT or *BAF155cKO* brains were electroporated with a *KAT2A*-expression plasmid. Compared with control (GFP-injected cortex) at E15.5, overexpression of *KAT2A* increased the percentage of proliferating BPs (GFP+/KI67+/TBR2+ or PAX6+) among targeted BPs (GFP+/TBR2+ or PAX6+) in WT (fig. S10G) and BAF155cKO cortex (fig. S10H). Furthermore, five out of six BAF155cKO and three out of six WT cortices injected with *KAT2A* expression plasmids at E12.5 displayed focal neocortex folding at E17.5 (Fig. 7H).

Collectively, these findings suggest that increased H3 acetylation at the promoter region of the evolution-related gene, TRNP1, selectively promotes the proliferative capacity of BPs, leading to enhanced neuronal output, increased cortical expansion, and gyrification during mammalian evolution (Fig. 7I).

## DISCUSSION

In this study, we describe H3 acetylation as a novel epigenetic mechanism that regulates neocortical expansion. We present evidence that human BPs have higher H3 acetylation levels than mouse BPs, and that elevated levels of H3ac preferentially promote BP self-amplification and augment neuronal output, leading to enlarged size and folding of the mouse neocortex (summarized in Fig. 7I). Mechanistically, this process involves the epigenetic and gene expression controls of the evolutionarily regulated gene, TRNP1, in cortical development.

### Contribution of BP proliferation to cortical expansion and folding

In lissencephalic rodents, APs in the VZ and TBR2+ BPs in the SVZ are largely responsible for neuron production during cortical development. Previous works showed that β-catenin overexpression-induced VZ progenitor amplification in the mouse brain yielded folding of the ventricular surface, not the cortical surface (*38*), and that increasing the pool of BPs/bIPs enlarged brain size but did not induce cortical folding (*10, 39*).

On the other hand, published evidence suggests that there is a link between the proportion of BPs/bRGs and the extent of cortical gyrification in various species (*25, 28, 40, 41*). bRG progenitors are derivatives of APs that presumably after becoming delaminated from their apical anchorage (Borrell and Gotz, 2014). However, when adherens junction proteins are lost (*31*) or the linkages to cytoskeletal belts are broken by downregulation of the small GTPase, RhoA, apically anchored progenitor cells are delaminated without significant change in bRG generation or cortical folding. Likewise, our data and that of others indicate that PAX6 and BAF155 control the delamination of RGs at least partly by regulating the expressional programs of genes encoding adherens junction proteins and RhoA (*31*). Similar to many other mouse mutants that exhibit delamination of APs during corticogenesis, BAF155cKO (*31, 42*) and PAX6cKO mice (*39, 43*) do not exhibit an enlarged and folded cortex, suggesting that the ectopic (outside of the VZ) localization of RGs may be insufficient to trigger cortical expansion and/or folding. Recent observations in the cortex of marmoset (a lissencephalic species) or agouti (a rodent with a moderately gyrencephalic cortex) revealed that the presence of an augmented population of bRGs in the expanded oSVZ of some mammalian subclasses may not correlate with the occurrence of cortical folding and gyrencephaly (*44, 45*). In gyrencephalic ferrets, for example, the oSVZ initially appears as a massing of numerous bRGs that are directly produced by the transient delamination of APs during early cortical development (*22*). Later, however, the generation of bRGs becomes completely independent of AP delamination, instead relying on their self-expansion (*22*). Along the same lines, functional blockade of TBR2 in ferrets was shown to cause premature neuronal differentiation of SVZ progenitors, diminishing the numbers of bIPs and bRGs and impairing gyrification (*46*). These findings support the idea that BP proliferation is a key driver of cortical folding (*46*). Thus, the simple delamination of VZ progenitors is insufficient to generate BPs with a high proliferative capacity; instead, this process seems to critically involve altered expression of regulatory factors that boost BP amplification.

### H3 acetylation promotes BP proliferation and cortical expansion

BPs from rodents and primates have distinct characteristics. In the lissencephalic rodent cortex, BPs are mainly neurogenic progenitors that express the transcription factor (TF), TBR2. In gyrencephalic species (such as ferret, primate, and human), in contrast, most of the BPs are proliferative progenitors and almost half of them co-express the TFs, TBR2 and PAX6 (*2*). In the developing ferret cortex, TBR2 is essential for the generation of both types of BPs (bIPs and bRGs) (*46*). A prerequisite for identifying the gene expression and epigenetic programs of BPs during evolution is thus the analysis of BP-specific signatures in distinct species. Recent single-cell transcriptomic and genetic studies have sought to elucidate the phylogenic expansion and gyrification of the mammalian cortex (*12–14, 32, 47–49*). A few factors (such as TRNP1, ARHGAP11B, and TBC1D3) have been found to control BP genesis, BP proliferation, cortical expansion, and folding (*10, 11, 13, 15–18, 39*). Recently, remodeling of chromatin via histone post-translational modifications has also been implicated in cortical neurogenesis. However, the epigenetic mechanisms that coordinate the expression or repression of genes that are essential for the developmental events underlying cortical expansion and folding during evolution are still largely unknown.

In this study, we purified TBR2+ BPs from mouse and human cortices. To compare the bulk level of epigenetic marks between mouse and human BPs, we quantified the PTMs of histones by simultaneously measuring the relative abundance of most known histone methylation and acetylation marks. This method requires a moderate number of cells that can be obtained by FACS, and allows for the systematic screening of epigenetic changes during evolution. We identified a few epigenetic marks that display different levels in TBR2+ BPs isolated from mouse versus human cortex. Given that PAX6+, SOX2+ BPs/bRGs are relatively rare in the WT mouse cortex (*25, 26*), we used the BAF155cKO mutant as a mouse model to investigate the effect of HDAC inhibition on the proliferation of BPs. Based on the expression of PAX6, AP2γ, and SOX2 in the VZ and pHH3 at the apical VZ surface (Fig. 4A/C), our data suggest that the elevation of H3ac in developing mouse cortex did not influence the proliferation and delamination of APs, which are consistent with a relatively high basal level of H3K9ac in both mouse and human APs. Intriguingly, we found that the level of H3K9ac is higher in human BPs than in mouse BPs. Moreover, we found that elevated H3K9ac specifically expands the pool of BPs, but not APs, and increases the number of both lower-layer CTIP2+ neurons and upper-layer SATB2+ neurons.

Cortical gyrification is underlined by the production of a large population of neurons and the presence of sufficient and more complex neuronal migration scaffolds, which permit lateral dispersion of neurons to form intricate cortical layers. In gyrencephalic species, cortical expansion is achieved by an extraordinary increase in neurogenic bIPs and the neurogenic/scaffold-forming bRGs in the oSVZ, which are vital in directing lateral dispersion of radially migrating neurons (*1–4*). The generation of more bIPs and bRGs in TSA-treated BAF155cKO cortex than in TSA-treated WT cortex could explain the corresponding differences in cortical size (Fig. 7A‒F) and level of cortical folding (fig. S10B) between these two groups. In agreement with previous studies (*19, 44, 45*), our findings suggest that the formation of cortical folding requires a great abundance of both bIPs and bRGs.

Cortical folding can be orchestrated by several genes known to critically regulate neuronal migration (*6*). For instance, the cell-adhesion molecules FLRT1/3 are crucial in directing cortical neuron radial migration, such that their deficiency caused cortical folds in mouse (*17*). In humans, defective reelin signalling caused by mutations severely impair cortical folding (*50*). In TSA-treated cortex (Fig. 7), the sulci correspond to ectopic reelin+ cells (Fig. 7C). These ectopic Cajal-Retzius neurons could promote aberrant trajectories of RG cells and abnormal migration of neurons. Although, our findings highlight a novel epigenetic mechanism underlying cortical folding possibly by increased proliferation of BPs, we do not exclude the possibility that H3 acetylation induces gyrification partly by atypical migration of neurons.

Notably, HDAC inhibition did not activate the expression of known human bRG genes such as HOPX, FAM107A. Our gene expression comparison revealed that TSA treated mouse BPs have closer molecular identity with macaque BPs than with those from human. In addition, an increased number of BP cells upon TSA treatment is not observed in the medial cortex. Apart from H3K9ac, the levels of several other epigenetic marks appeared to be higher in human BPs at GW14 and GW18 than in mouse BPs at E13.5 and E16.5 (Fig. 1D), including H3K18ac, H3K4ac, and H3K4me2, which were also previously shown to be enriched at promoters and enhancers, and to activate transcription. These findings suggest that the high proliferative capacity of BPs in medial cortex and gene expression program of BPs requires high levels of several other epigenetic marks and not on H3K9ac exclusively. Our on-going work aims to examine whether increased H3K4me2 in cooperative action with H3K9ac could promote expression of BP genes and is able to induce proliferation of BPs in medial cortex.

Interestingly, our GO analysis revealed a relatively high expression of genes coding for histone deacetylases (such as HDAC1, HDAC2, HDAC3, HDAC8, HDAC9, fig. S2F-H) in TBR2+ BPs in mice. This could explain why levels of H3ac are low in mouse BPs. It will therefore be particularly interesting in the future to investigate the upstream mechanisms and analyze how differential H3 acetylation across different species is modulated.

Together, our findings indicate that H3 acetylation has prominent impacts on the cerebral cortex, in terms of both radial and tangential expansion (Fig. 7I), by increasing the population of highly proliferative BPs and thus the degree of cortical gyrification (as seen in the primate brain).

### TRNP1 is a main target gene of H3 acetylation in BP amplification

Our RNA-Seq and H3K9ac ChIP-Seq analyses for many genes in the sorted TBR2+ BPs indicate that HDAC inhibition either increases the H3K9ac level or upregulates the expression of the studied genes. Interestingly, the correlation of these data indicates that H3K9ac directly activates the expression of only a few genes (Fig. 5C-H). Such genes include TRNP1, whose manipulated expression in the mouse and ferret cerebral cortex was previously reported to affect various cortical expansion-related features, including the proliferation of cortical neuroprogenitors and the induction of cortical folding (*11, 22, 51*). We herein show that H3K9ac positively regulates the expression of TRNP1 in BPs to orchestrate BP proliferation in the developing cortex. Expressional analyses indicate that the expression patterns of H3K9ac (this study) and TRNP1 (*11*) are similar in developing mouse and human cortex. In particular, H3K9ac and TRNP1 exhibit low or undetectable levels in mouse BPs, whereas their expression is much higher in human BPs. Furthermore, we show that the overexpression of TRNP1 by means of H3K9ac epigenome editing increases BP proliferation, as also seen in HDAC inhibitor-treated cortex, and that this phenotype is largely reverted to control levels upon TRNP1 knockdown. These findings bring to light the role of H3 acetylation as a mechanism upstream of TRNP1 function for BP proliferation in cortical development during evolution.

In our observation, the expression of H3K9ac-direct targets such as ADRB1, TRNP1 and PCDH1 is very sensitive and relatively proportional to the level of H3K9ac at their promoters. The TSA treatment led to about 1.5–2.3-fold increase in H3K9ac level at the promoter region and about 2.6–4.0-fold increase in expression of these genes (Fig. 5D/H). In the same line of evidence, the treatment of gTRNP1#2 increased 1.20-fold in H3K9ac level at its promoter and 1.23-fold in its expression level (Fig. S9H/I). Remarkably, the moderately-increased level of H3K9ac at the promoter and expression of key proliferative factor TRNP1 (*11, 22*) by the gTRNP1 electroporation is sufficient to evoke the proliferation of TBR2+ BPs in developing mouse cortex (Fig. 6D/E/H). Our findings also suggest that the increased expression of many genes upon the TSA treatment might synthetically promote the proliferation of BPs. It is worth determining the function of ADRB1, and PCDH1 alone or together with TRNP1 in BP amplification and cortical expansion.

In summary, we herein identify a mechanism whereby BP amplification is epigenetically controlled through H3 acetylation, which specifically affects the expression level of TRNP1. Our findings show that the temporally dynamic regulation of H3 acetylation and its downstream effector, TRNP1, are part of an evolutionary epigenetic mechanism aimed at enlarging the diversity of neocortical phenotypes during evolution.

## MATERIALS AND METHODS

### Plasmids

Plasmids used in this study: pCIG2-KAT2A-ires-eGFP, gTRNP1-dCas9-KAT2A-T2A-eGFP (this study), shTRNP1 constructs (*11*).

### Antibodies

The following polyclonal (pAb) and monoclonal (mAb) primary antibodies used in this study were obtained from the indicated commercial sources: H3K9Ac rabbit pAb (Abcam), SOX2 mouse mAb (1:100; R&D Systems), SOX2 goat (1:100; Santa Cruz), PAX6 mouse mAb (1:100; Developmental Studies Hybridoma Bank), PAX6 rabbit pAb (1:200; Covance), TBR2 rabbit pAb (1:200; Abcam), CidU rat pAb (1:100; Accurate), KI67 rabbit pAb (1:50; Vector), GFP rabbit pAb (1:1000; Abcam), GFP chick pAb (1:1000; Abcam), AP2γ mouse mAb (1:100; Abcam), phospho-H3 rat pAb (1:300; Abcam), pVIM mouse mAb (1:500; MBL), TNC rabbit pAb (Abcam), PTPRZ1 rabbit pAb (Sigma), BAF170 rabbit pAb (Bethyl), BAF170 rabbit pAb (Sigma), BAF155 rabbit pAb (1:20; Santa Cruz), BAF155 mouse mAb (1:100; Santa Cruz), CASP3 rabbit pAb (1:100; Cell Signaling), CTIP2 rat pAb (1:200; Abcam), HuCD mouse mAb (1:20; Invitrogen), TBR1 rabbit pAb (1:300; Chemicon), SATB2 mouse mAb (1:200; Abcam), CUX1 rabbit pAb (1:100; Santa Cruz), β-actin rabbit pAb (Sigma), H3ac rabbit pAb (Upstate), NeuN mouse mAb (Chemicon), REELIN mouse mAb (Gift from Prof. Goffine), SOX5 rabbit pAb (Santa Cruz), KAT2A rabbit pAb (Abcam), NESTIN mouse mAb (BD), RC2 mouse mAb (Developmental Studies Hybridoma Bank),

Secondary antibodies used were horseradish peroxidase (HRP)-conjugated goat anti-rabbit IgG (1:10000; Covance), HRP-conjugated goat anti-mouse IgG (1:5000; Covance), HRP-conjugated goat anti-rat IgG (1:10000; Covance), and Alexa 488-, Alexa 568-, Alexa 594- and Alexa 647-conjugated IgG (various species, 1:400; Molecular Probes).

### Human fetal brain collection and processing

Human fetal brain tissue was obtained from spontaneous abortions that occurred in the Hospital for Obstetrics and Gynecology “Prof. Dimitar Stamatov”, Medical University – Varna, Bulgaria, after an informed written maternal consent, and with the approval of the local Ethics Committee (Protocol No 19/April 2012; Protocol No 55/June 2016) according to IRB guidelines. The gestation age in weeks (GW) has been determined based on the history of last menstruation as reported by the patients. The heads were placed in 4% PFA solution in PBS 7.5 pH for 24h, and then the brains were dissected and postfixed in PFA for 5-7 days, cryoprotected in sucrose and frozen in OCT medium prior to cryosectioning at 20 μm.

Cortical tissues from additional cases were used for FACS as described for mouse tissues in the below section.

### Animal care, generation of transgenic mice, *in utero* electroporation

Floxed BAF155 (*52*), Emx1-Cre (*53*) mice were maintained in a C57BL6/J background. *In utero* electroporation was performed as described previously (*42, 54*). Animals were handled in accordance with the German Animal Protection Law and with the permission of the Bezirksregierung Braunschweig according to IACUC guidelines.

### Mouse treatment with HDAC inhibitors (HDACi)

Trichostatin A (TSA) (Sigma-Aldrich, Cat. T8552-1MG) was dissolved in vehicle (8% ethanol in 1xPBS) at a concentration of 100µg/ml. Suberoylanilide hydroxamic acid (SAHA) (Biomol, Cat. CAS 149647-78-9) was dissolved in Vehicle (DMSO) (10 mg/ml). Sodium salt Valproic acid (VPA) (Sigma, Cat. P4543) was dissolved in Vehicle (saline) at concentration of 100 mg/ml. Pregnant mice from E12.5 d.p.c. were injected intraperitoneally twice daily (except the experiment to examine a dose-dependent effect in fig. S4A/B) with either vehicle or 150µl of 100 µg/ml TSA solution or 20µl of 10 mg/ml SAHA solution plus 110 µl Saline or 120µl of 100 mg/ml VPA solution. Treated mice were sacrificed at different developmental stages as indicated in the text.

### TBR2+ nuclei and cell sorting from embryonic cortex

Since typical cell sorting methods comprise of protease treatment for cell dissociation that can lead to unwanted effects on gene expression profile, we opted for unbiased approach to minimize biasness due to sample processing. In this regard, we opted for TBR2+ nuclei sorting from fresh mouse cortex, instead of cell sorting, as this is a well-established method of getting the cell types of interest from the brain without the biasness of the sample preparation (*23*). Since we were interested to obtain epigenetic and transcriptomic data from TBR2+ and TBR2-nuclei, two different protocols were followed, respectively. By unknown reasons, TBR2 antibody did not work in our intracellular immunostaining protocols for nuclei, which were prepared from frozen tissue.

In our mass spectrometry analysis, we used frozen human cortex as the cell source for sorting. We therefore establish a cell preparation protocol using solution with stringent reagents for cell sorting.

#### TBR2+ nuclei sorting protocol from embryonic mouse brain for ChIP-Seq

The protocol were adapted originally from a previously published protocol(Halder et al., 2016). All steps were done on ice or at 4 degrees unless stated otherwise. Freshly prepared embryonic cortices from 5 CD1 pups were pooled for each replicates and homogenized briefly in low-sucrose buffer (320 mM Surcrose, 5mM CaCl2, 5 mM MgAc2, 0.1 mM Ethylenediamine tetraacetic acid (EDTA), 10mM HEPES pH 8, 0.1% Triton X-100, 1mM DTT, supplemented with Roche protease inhibitor cocktail) with plastic pestles in 1.5mL tubes. Crosslinking was done with 1% Formaldehyde and incubated for 10 minutes at room temperature on a rotating wheel. 125mM Glycine was used for quenching remaining formaldehyde in solution by incubation for 5 minutes at room temperature. After centrifugation for 2000xg for 3 minutes, the crude nuclear pellet was pipette mixed into additional low-sucrose buffer and further homogenized with a mechanical homogenizer (IKA Ultraturax). The solution was carefully layered on high-sucrose buffer (1000mM Sucrose, 3mM Magnesium acetate, 10mM HEPES pH 8, 1 mM DTT, protease inhibitor) in oak-ridge tubes and spinned at 3200xg for 10 minutes in a swinging bucket rotor centrifuge. This was done to get rid of myelin. Supernatant was carefully removed without disturbing the nuclear pellet and transferred into 2mL microfuge tubes(DNA-low bind). Nuclei were collected by centrifuging for 3 minutes at 2000xg. After removing remaining sucrose buffer, nuclei were resuspended into 500uL PBTB buffer(<u>PB</u>S-0.2% <u>T</u>ween-1% <u>B</u>SA buffer, with protease inhibitor). Tbr2 staining was performed using Anti-TBR2(Eomes)-Alexa488 conjugated antibody(IC8889G-025, 1:100) for 1 hour. Nuclei were washed once and finally resuspened into 500µl PBTB. FACSAria III with was used to performing Tbr2 nuclei sorting, while samples without antibody acted as negative control for gating. Nuclei were sorted directly into PBTB coated 15mL falcon tubes and centrifuged briefly to pellet the Tbr2 positive and negative nuclei. Sorted nuclei ellets were flash frozen into liquid nitrogen and stored at −80°C, before proceeding with ChIP experiment.

#### TBR2+ nuclei sorting protocol from embryonic mouse brain for RNA isolation and RNA-Seq

Freshly dissected mouse embryonic cortices from 5 CD1 pups were pooled and immediately submerged into enough RNAlater solution in a microfuge tube and kept at 4°C for at least 24 hours. The excess RNAlater solution were pipetted out and washed twice with 1x RNAse free PBS. After last wash, all the incubation steps were done on ice or centrifugation at 500xg at 4° unless stated otherwise. The tissues were immediately submerged into 500 uL lysis buffer (Sigma, NUC101), dounce homogenized using plastic pestles for 30-45 times and additional lysis buffer was added to make the volume up to 2 mL. After incubating for 7 minutes, lysates were centrifuged for 5 minutes and nuclei pellet was resuspended into 2mL lysis buffer. Lysates were incubated for 7 minutes, filtered through 40µm filter into a new 2mL tube and centrifuged to remove supernatant. Pellet was washed with 1800uL nuclei suspension buffer (NSB; 0.5% RNAse-free BSA(Millipore), 1:200 RNaseIN plus RNAse inhibitor(Promega), protease inhibitor diluted into RNAse free PBS(Invitrogen)) by centrifuging and resuspended in 500uL NSB. Tbr2 staining was done using Anti-TBR2(Eomes)-Alexa488 conjugated antibody(IC8889G-025, 1:100) for 1 hour. Stained nuclei were washed and resuspended both using NSB. Nuclei sorting were performed as aforementioned section(Tbr2 nuclei sorting for ChIPseq) while sorted nuclei were collected into NSB coated falcon tubes. Nuclei were collected with brief centrifugation and Trizol LS solution was added for RNA isolation. After adding chloroform and centrifugation for 15 minutes at 120000xg, aqueous phase was collected and RNA was purified with Zymo RNA clean & concentrator-5 kit. Resulting RNA was used with Takara SMART-Seq v4 Ultra Low Input RNA kit to prepare mRNA-seq libraries using 1ng of RNA from sorted Tbr2 nuclei. Libraries were sequenced in Illumina Hiseq 2000 machine.

#### TBR2+ cell sorting protocol from embryonic mouse brain for mass spectrometry analysis

Pool at least 8 (at E13.5) or 4 (E16.5) mouse cortices or equal amount of human cortex per replicate. Cortical cells were dissociated by using the Neural tissue dissociation kit (MACS Miltenyi Biotec, #130-092-628 P). Subsequently, cells were fixed and permeabilized with the Foxp3 Fixation/Permeabilization working solution (Foxp3/Transcription Factor Staining Buffer Set, eBioscience, #00-5523). For antibody labeling, cells were incubated with Alex 488 conjugated TBR2 antibody (1:200 in Permeabilization Buffer) for at least 30 min on ice (in the dark) with slight shaking. Stained cells then were washed twice with Permeabilization Buffer. After sorting, pellet of the sorted cells was collected by centrifugation. Samples should be stored in −80°C if the experiment are not done immediately.

### Mass Spectrometry analysis of epigenetic marks

Histones were prepared from 2.5×10^6^ sorted TBR2+ cells for each MS sample. Histone extraction, mass spectrometry (MS) analysis were performed as described previously (*55*). To generate the heatmaps, for each individual modification, the data for each sample was converted to the fraction of the sum across all samples to display relative abundances across the sample group, and then conditionally formatted in Excel using the default red/white/blue color scheme. The hierarchical clustering heatmap was generated in R using the “pheatmap” function with default settings.

### Chromatin immunoprecipitation (ChIP)

The ChIP protocol was described previously (*56*) with H3K9ac antibody (Millipore 07-352). Resulting ChIPed DNA was used for either qPCR or sequencing for appropriate experiments. For ChIP sequencing, immunoprecipitated DNA was further processed with NEBNext Ultra II DNA library preparation kit to generate Illumina sequencing libraries. Libraries were sequenced in Illumina Hiseq 2000 machine

### Next generation sequencing data analysis (ChIP-seq, RNA-seq)

Base calling and conversion to fastq format were performed using Illumina pipeline scripts. Afterwards, quality control on raw data was conducted for each library (FastQC, www.bioinformatics.babraham.ac.uk/projects/fastqc). The following control measurements and information were obtained: per base sequence quality, per sequence quality scores, per base sequence content, per base GC content, per sequence GC content, per base N content, sequence length distribution, sequence duplication levels, overrepresented sequences, Kmer content. The reads were mapped to a mouse reference genome (mm10) using bowtie2. *rmdup* function of samtools (*57*) was used to remove PCR duplicates from each BAM file. *merge* function of samtools was used for merging BAM files with unique reads (i.e., with duplicates removed) belonging to replicates from the same group into a single BAM file. All the downstream analyses were performed on BAM files with only unique reads. Profile plots of H3K9ac were created with NGSPlot using merged BAM files from immunoprecipitated samples and inputs. H3K9ac enrichment at different gene loci was visualized through the Integrated Genome Browser using wiggle files that were created from the merged BAM files with the script from the MEDIPS package of Bioconductor. Peaks were called on individual with MACS2 with q-value < 0.1. Differential binding was assessed with DiffBind package of Bioconductor (*58*) with in-built DESEQ2 option implemented in differential analysis. In differential binding analyses promoters were defined as +/- 2000 bp from transcription start site (TSS). Peak annotation was done with HOMER. RNA-seq was performed as described previously (*56, 59*).

### Epigenome editing

#### Constructs for epigenome editing

The backbone vector encoding sgRNAs and 3xFlag-dCas9 (nuclease dead-Cas9)-T2A-GFP expression cassette was generated from the vector pSpCas9n(BB)-2A-GFP (PX461 in addgene): H840A was introduced into Cas9n to produce dCas9 through PCR-mediated mutagenesis cloning, and a polylinker sequence 5’-TCCGGACGGGGATCCACTAGTGTCGACACCGGTCCTAGG-3’ was inserted right after the Cas9 coding sequence. The vectors for the mouse TRNP1 promoter epigenome editing were derived from the above generated backbone vector. gRNA sequences targeting specific mouse TRNP1 promoter region (Chr4: 133497514-133498529) was designed using the software ‘Genious’, and cloned into the vector between BbsI sites after in vitro test, then the coding sequence for mouse KAT2A was amplified from the vector pCMV-sport2-mGCN5 (gift from Sharon Dent, Addgene plasmid # 23098), and cloned into the backbone vector between SpeI and AvrII sites, in frame with that of dsCas9 and T2A-GFP. The vectors were confirmed by sequencing.

#### In vitro test for TRNP1 sgRNAs, quantification of H3K9ac level at TRNP1 promoter and expression of TRNP1 in gTRNP1-dCas9-KAT2A-transfected Neuro2A

For in vitro test, sgRNAs were synthesized through in vitro transcription from a PCR-produced sgRNA template, which habors the T7 promoter abided by the gRNA sequence and the sharped gRNA scaffold using MEGAscript T7 Transcription Kit (Invitrogen) according the manufacturer’s protocols. Cas9 protein was purchased from IDT (#1074182). In vitro test of Cas9/gRNAs complexes cutting efficiency was performed using a 1016 bp PCR product from TRNP1 promoter according the protocol described in (*35*).

Based on the in vitro testing result, the gRNA2 and gRNA4 were selected and cloned into above mentioned vector to generate vectors TRNP1-sg2-Flag-dCas9-2A-GFP (TRNP1-sg2), and TRNP1-sg4-Flag-dCas9-2A-GFP (TRNP1-sg4). 12 µg of parental vector Flag-dCas9-2A-GFP or TRNP1-sg2, or/and TRNP1-sg4 were transfected into Neuro2A cells cultured on 10 cm dishes using lipofectamine 2000 reagent (thermofisher) following the manufacture’s protocol, and cells were harvested for FACS analysis 3 days post-transfection for qPCR and ChIP/qPCR analyses.

### Cell cultures

For cell culture assays, plasmids were transfected into Neuro2A cells using Lipofectamine 2000, or were electroporated into primary cortical cells using a mouse neural cell Nucleofector kit and a nucleotransfection device (Amaxa).

### IHC, cell-cycle parameter and DiI labeling experiments

IHC. determination of cell-cycle index and DiI labeling were performed as previously described (*35, 43*).

### Retrovirus production and viral infection in slices

Retrovirus for GFP labeling was produced from 293gp NIT-GFP packaging cells by transfection with pCMV-VSV-G plasmid (Addgene #8454). Viral particles with low titer of about 10^6^ PFU/ml were used for infection. For the viral infection, cortical slices from brain of TSA-treated BAF155cKO embryos were prepared and cultured as described previously (*56*). To label the bRGs, GFP-expressing retrovirus (106 PFU/ml) was injected into IZ of the cultured cortical slice, using a glass micropipette. 60 hours post-infection, brain slides were washed and stained with antibodies against GFP and PAX6.

### qRT-PCR and WB analyses

qRT-PCR and WB analyses were performed as described previously (*54*) using primers described in Supplementary Table S7 and the RT² Profiler PCR Array profiles (Qiagen).

### Relative quantification of cortical surface, cell counts and quantitative analysis of immunohistochemical signal intensity

The cortical surface was measured and analysed as described previously (*31, 39*). For the quantitative analyses of immunohistochemical signal intensity of H3K9ac and H3ac, confocal fluorescent images of sections of mouse and human cortex were used. The IHC Images with corresponding IgG isotopes were used as immune-staining controls. The color images of cortex were converted to gray scale to eliminate background. The pixel values of the fluorescent signal intensity were measured by using Analyze/Analyze Particles function (ImageJ software) as previously described (*39, 60*). The measured value was then subtracted from immune-staining controls (without primary antibody).

Similarly the relative amount of protein from developed films in WB experiment was quantified densitometrically using ImageJ software as described previously (*39, 60*).

Immunomicrographs were quantified using anatomically matched forebrain sections from control and mutants or HDACi-treated embryos. Marker-positive cells within images of the cortex obtained by confocal microscopy were counted for comparison.

In developing mouse cortex, expression level of Pax6 is very variable from Pax6^low^ to Pax6^high^ in cortical progenitors (*54*). In some cells, expression of Pax6 is even at the border between background and very low (Pax6^very low^). We noted that TSA treatment massively increased the number of Pax6+ cells in SVZ/IZ. Among these Pax6+ cells, many are Pax6^very low^. In our cell counting and quantification for Pax6+ cells, images were acquired with confocal microscopes and were further analyzed with Adobe Photoshop. We adjusted the brightness of images for both TSA- and Veh-treated samples to mask Pax6^very low^ cells.

Generally, cell counts of six matched sections were averaged from three biological replicates. Student’s *t*-test was used to statistically analyze histological data. All statistical tests are two-tailed, and *P*-values are considered significant for α = 0.05. Bar graphs are plotted as means ± SEM. Details of statistical analyses of histological experiments are presented in Table S9.

### Image acquisition and statistical analysis

Images were acquired with epifluorescence (Leica DM 6000) and confocal microscopes (Leica TCS SP5). Adobe Photoshop was used to process images for further analyses. Statistical significance was determined by Student’s *t*-test or Mann-Whitney test. All graphs are plotted as mean ± SEM. The statistical quantification was performed as average from at least three biological replicate. All details of statistical analyses for histological experiments are presented in Table S9.

## ACKNOWLEDGMENTS

We acknowledge T. Huttanus and H. Fett, U. Kunze, C. Heuchel, U. Teichmann for their expert animal care. We thank K. Jones for providing reagents, and C. Dean for helpful discussions. This work was supported by the Research Program, Faculty of Medicine, Georg-August-University Goettingen (TT), TU432/1-1, TU432/3-1, TU432/6-1 DFG grants (TT), Schram-Stiftung (TT, AF), DFG-CNMPB (TT, JS, AS, AF), ChroNeuroRepair (MG) and advanced ERC grant (MG).

## AUTHOR CONTRIBUTIONS

CK performed ChIP-seq, RNA-Seq and the corresponding bioinformatics analyses; LP and TT performed most histological analyses of IUE and HDACi experiments; AT, LP, RM, EK, MA and TT collected human fetuses and performed analyses of expression patterns of mouse and human cortices; MSS and LK contributed to ChIP-Seq library preparation, establishment of TBR2+ cell sorting, and cell-type specific RNA-Seq. YX, GS, HN, JR, PAU and EA contributed in histological analyses, epigenome editing, and statistical quantification. AM contributed to qPCR and ChIP-qPCR. VC performed QC analysis and alignment of ChIP-Seq and RNA-Seq reads. MA and WH contributed in epigenome editing. ME and MG contributed to TRNP1 study. AS, HPN, RHS and JS provided transgenic lines and research tools; JFS, AS, WH, MG and AF offered suggestions for the study; TT conceived the study and wrote the draft. CK, GS, JFS, MA, ME, AT and AS revised the manuscript.

## DATA AVAILABILITY

All RNA-Seq and ChIP-Seq data have been deposited in GEO and will be released to public upon acceptance of the manuscript.

## Supplementary Materials

**Fig. S1.**
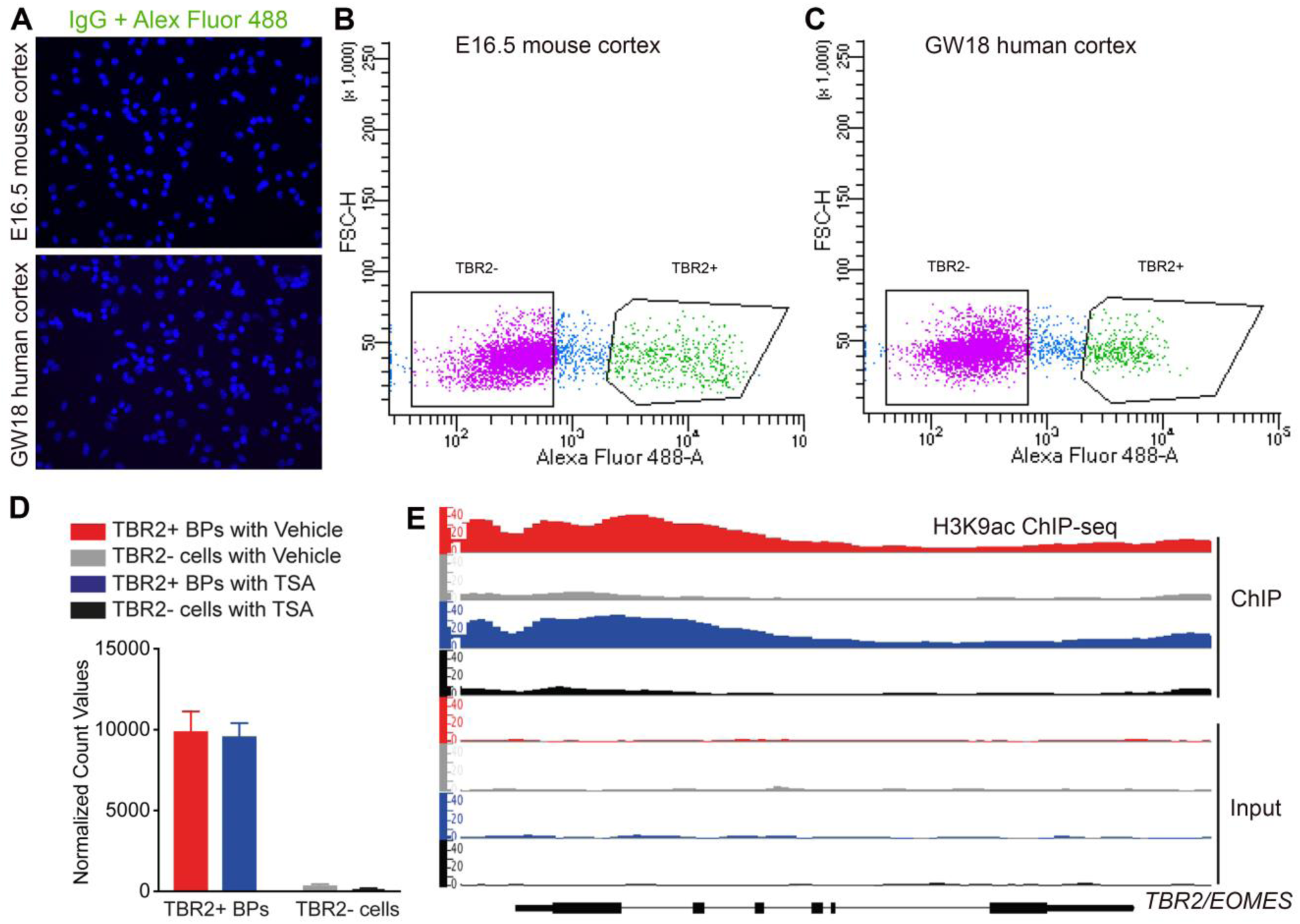
Purification and characterization of TBR2+ BPs. (A) Images showed stained samples as isotype control (see also Fig. 1B/C). (B, C) Representative plot showing sorting gates for TBR2+ cells and TBR2-cells from mouse cortex at E16.5 (B) and human cortex at GW18 (C). (D, E) Purity of the sorted TBR2+ BPs and TBR2-cells was further proven by RNA-Seq (D) and H3K9ac ChIP-Seq (E). TBR2+ BPs had much higher levels of TBR2 RNA expression (D) and much higher levels of H3K9ac at TBR2 gene locus (E) compared to TBR2-cells.

**Fig. S2.**
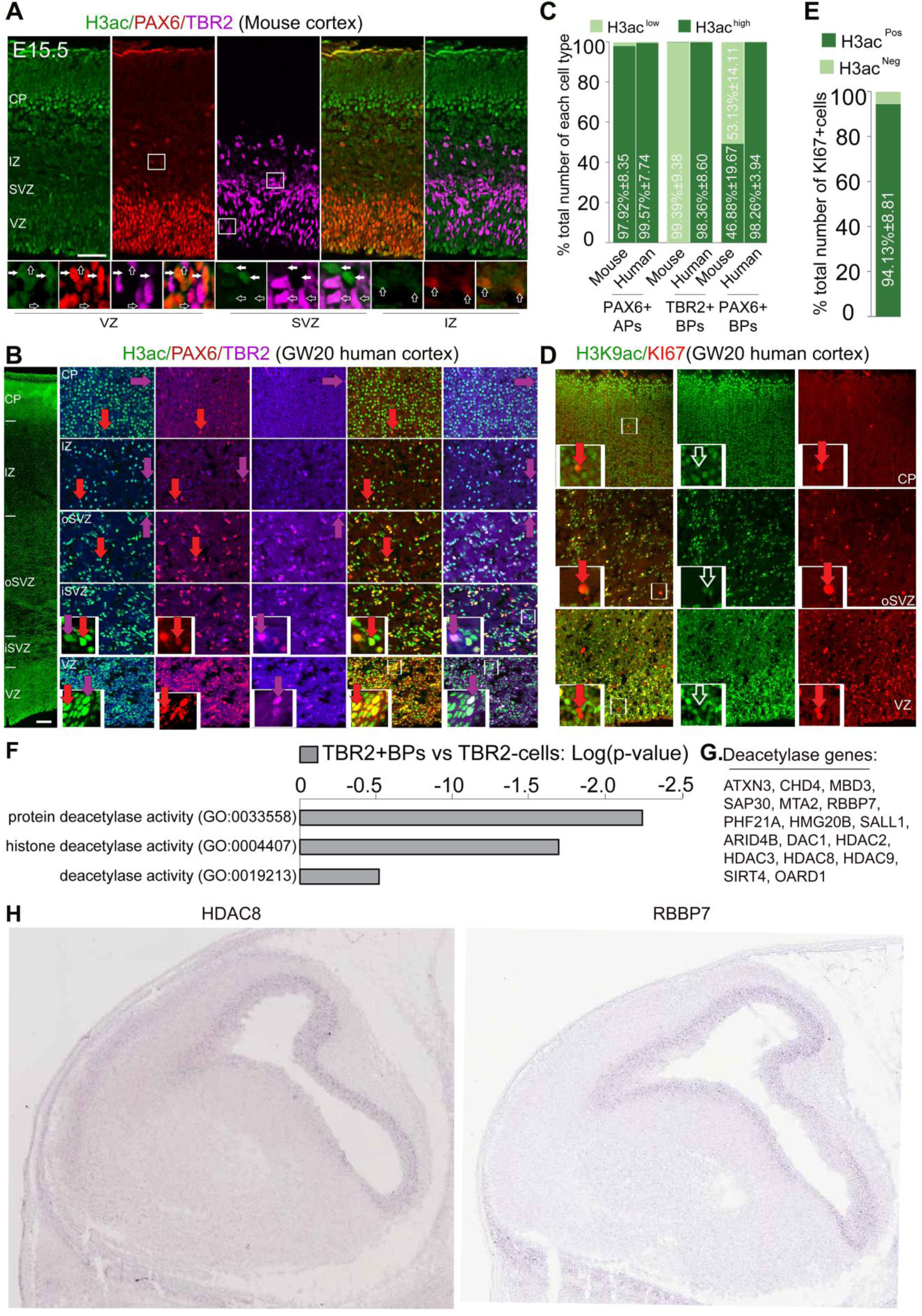
Histone H3 is acetylated differently in basal progenitors in murine and human developing cortex. (A) Images of triple IHC with cortical section from mouse embryo at E15.5 stained with antibodies against: H3ac, PAX6, and TBR2. The lower panels showing selected frames at VZ and IZ at higher magnification reveal PAX6^high+^, TBR2^low+^ APs are H3ac^high+^ cells (filled arrows), whereas many PAX6^high+^ BPs in IZ and all TBR2^high+^ BPs are H3ac^low+^ cells (empty arrows). (B) Images showing triple immunolabeling with antibodies against H3ac, PAX6 and TBR2 and human fetal cortex at GW20 demonstrate that both the PAX6+ APs and PAX6+ BPs (arrows in red) and the TBR2+ BPs (arrows in magenta) are highly immunoreactive with H3ac. (C) Statistical analyses of IHC assay (shown in A and B) comparing the level of H3ac (H3ac^high^ and H3ac^low^) in progenitor subtypes in developing mouse and human cortex confirm the high level of H3ac mark in both APs and BPs in humans. (D, E) IHC (D) and statistical analyses (E) of GW20 human cortical sections with H3K9ac and KI67 antibodies revealed that not all H3K9ac+ cells co-express KI67. (F) Graphical representation of Gene ontology analysis with terms related to protein deacetylases. (G) List of the genes identified in mouse BPs that functionally fall under protein/histone deacetylation processes. (H) Micrographs showing *in situ* hybridization obtained from GenePaint database, and indicating examples of the identified protein/histone deacetylation genes with distinctive expression in the developing mouse cortical subventricular zone. Abbreviations: BPs, Basal progenitors; vRGs, ventricular radial glial progenitors; IPs, intermediate progenitors; oRGs, outer sub-ventricular radial glial progenitors; VZ, ventricular zone; SVZ, subventricular zone; iSVZ, inner subventricular zone; oSVZ, outer subventricular zone; IZ, intermediate zone; CP, cortical plate. Scale bars = 50 μm.

**Fig. S3.**
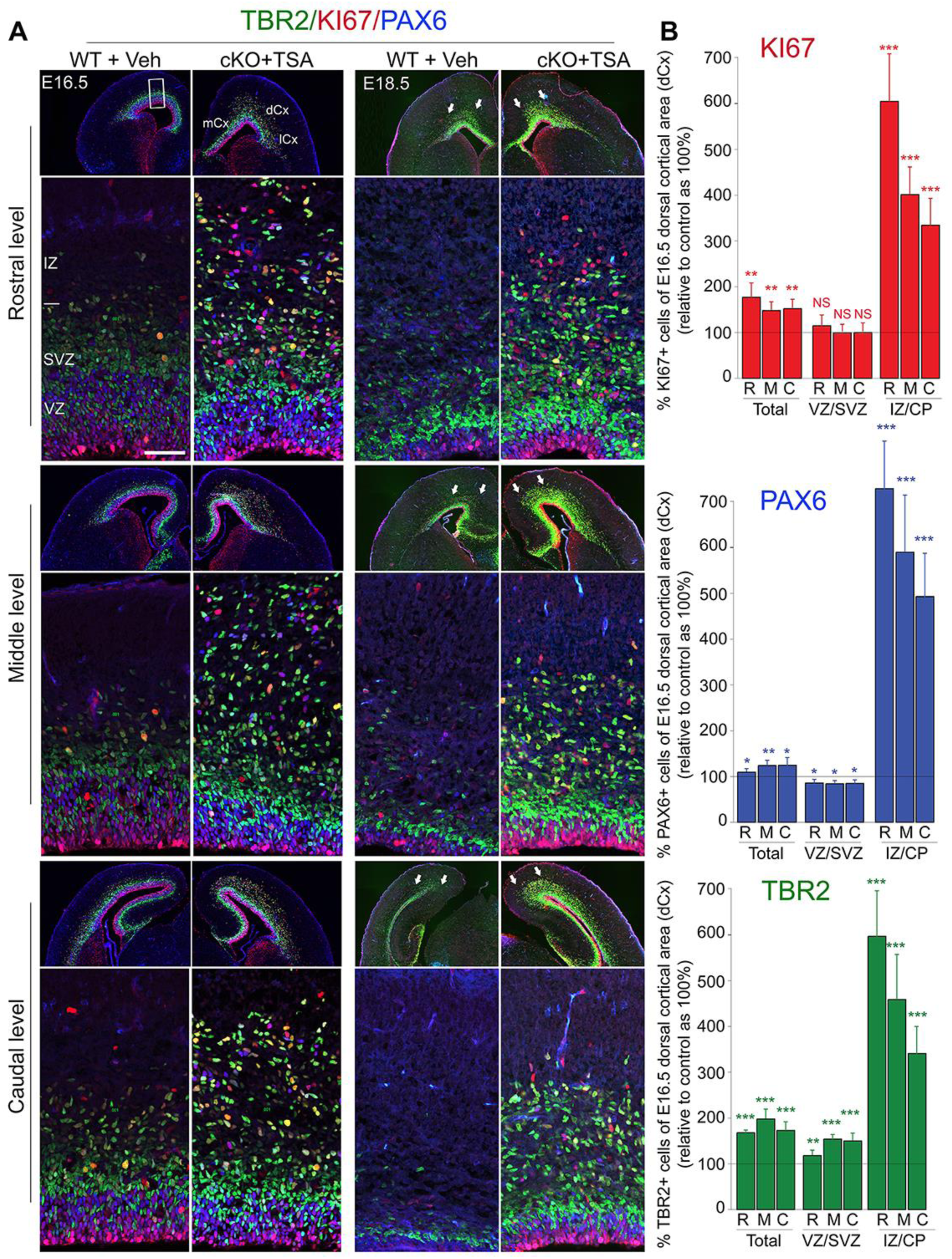
Area-restricted increases of TBR2+, PAX6+, KI67+ BPs in HDACi-treated cortex. (A) Merged confocal micrographs of fluorescent immunostaining for PAX6 (APs and BPs), TBR2 (BPs), and KI67 (actively-proliferating progenitors in cell cycle) with rostral, middle and caudal levels from unfolded areas of Veh-treated WT and TSA-treated BAF155cKO cortex at stages E16.5 and E18.5. In each level, *upper rows* present coronal mouse brain sections while *lower rows* present confocal images of an area in dorsal pallium. The arrows point the border between dCx/mCx, which corresponds to the selected medNcx region at E18.5 in the previous study (Vaid et al., 2018), and contains a high proportion of Tbr2+, Pax6+, Ki67+ BPs. (B) Statistical analyses of immunostainings shown in A for dorsal cortex (dCx) at E16.5. Values are presented as mean ± SEM (*P <0.05, **P <0.01, ***P <0.005). Abbreviations: BPs, Basal progenitors; vRGs, ventricular radial glial progenitors; IPs, intermediate progenitors; VZ, ventricular zone; SVZ, subventricular zone; IZ, intermediate zone; CP, cortical plate. Scale bars = 50 μm.

**Fig. S4.**
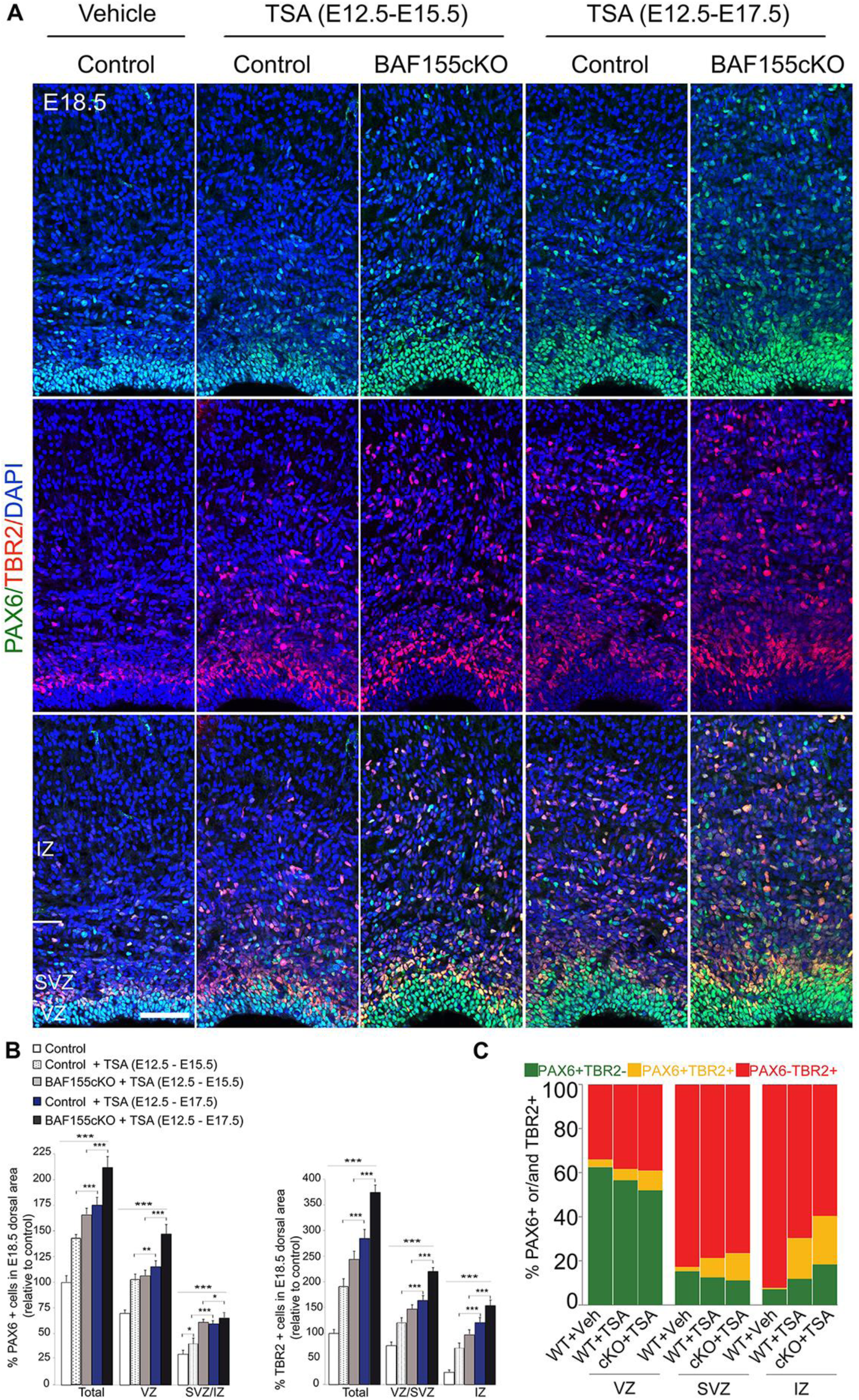
HDAC inhibition increase the genesis of BPs in dose-dependent fashion. (A) Immunofluorescence micrographs showing the quantity and distribution of PAX6 and TBR2 at E18.5 in control and BAF155cKO cortices, with or without two regimens of HDACi treatment. (B) Statistical analysis for dorsal cortical area revealed that more PAX6+ BPs and TBR2+ BPs were found in E18.5 cortex treated with TSA for E13.5-E17.5 than those in those injected with TSA for E13.5-E15.5. Notably, the TSA treatment also led to increased number of PAX6+vRGs at E18.5. (C) Proportions of cortical progenitors expressing PAX6 and/or TBR2 in VZ, SVZ, and IZ from dorsal area of WT+ Veh, WT+ TSA and BAF155cKO+ TSA cortex at E16.5. Values are presented as mean ± SEM (*P <0.05, **P <0.01, ***P <0.005). Abbreviations: BPs, Basal progenitors; vRGs, ventricular radial glial progenitors; IPs, intermediate progenitors; VZ, ventricular zone; SVZ, subventricular zone; IZ, intermediate zone; CP, cortical plate. Scale bars = 50 μm.

**Fig. S5.**
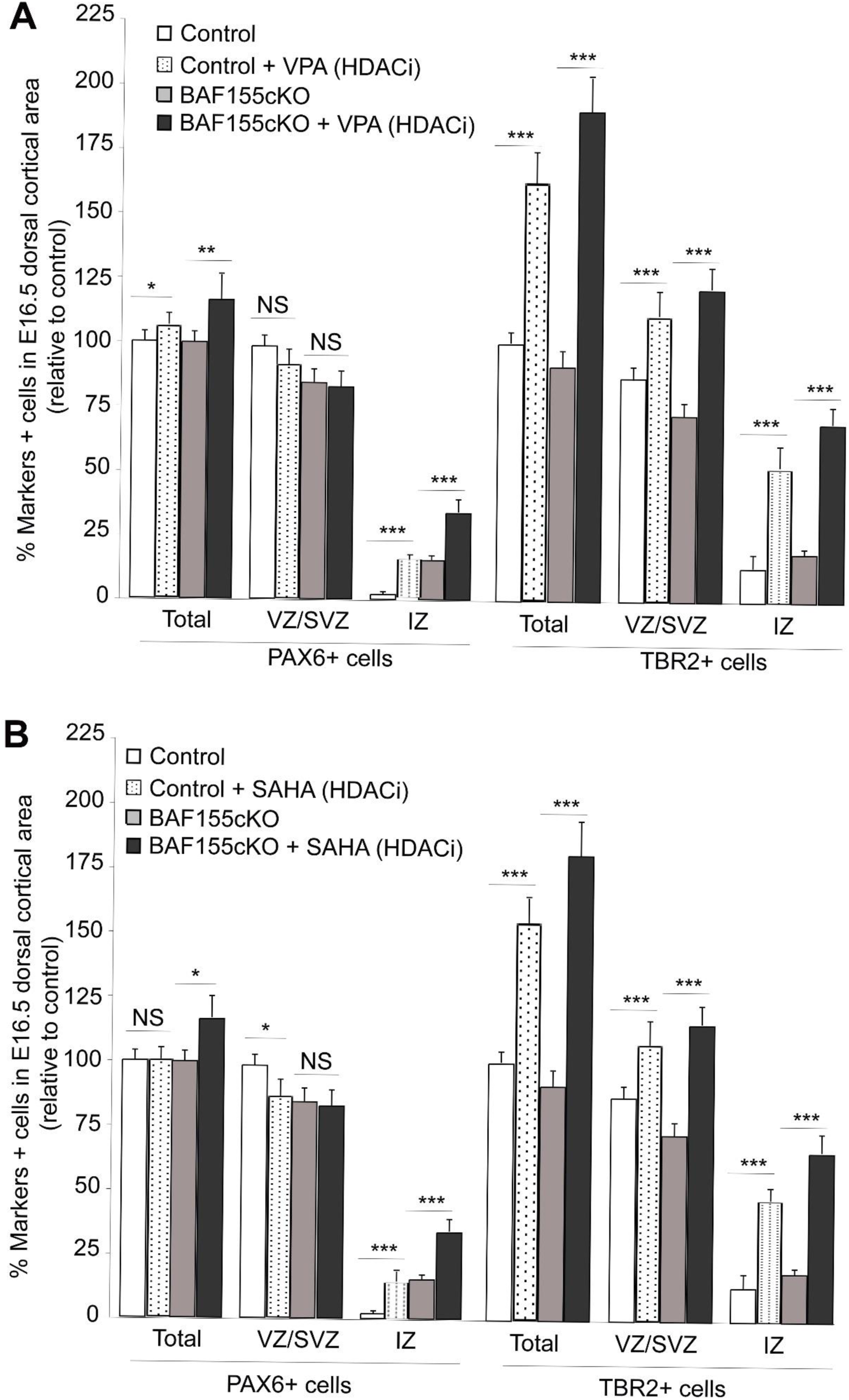
Increased level of H3ac promotes genesis and proliferation of BPs in developing mouse cortex. (A, B) Increasing H3 acetylation by other HDAC inhibitors such as VPA (A) and SAHA (B) also increased the number of TBR2+, PAX6+ BPs in both WT and BAF155cKO developing cortex. Values are presented as mean ± SEM (*P <0.05, **P <0.01, ***P <0.005). Abbreviations: BPs, Basal progenitors; vRGs, ventricular radial glial progenitors; IPs, intermediate progenitors; VZ, ventricular zone; SVZ, subventricular zone; IZ, intermediate zone; CP, cortical plate. Scale bars = 50 μm.

**Fig. S6.**
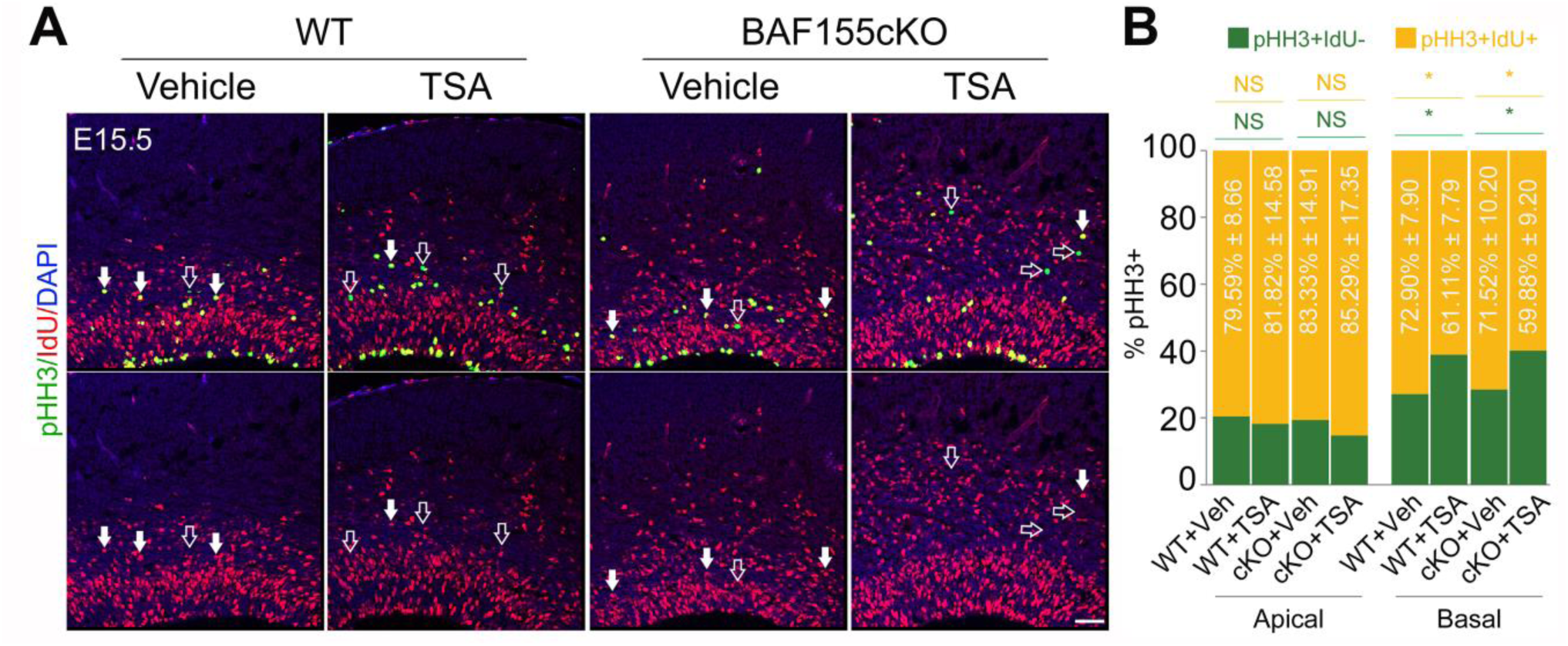
H3 acetylation specifically promotes proliferation of basal progenitors. (A, B) 4 hours-IdU pulse-labeling is to trace the progression of cortical progenitors within the S-M phases. (A) Double IHC analysis with antibodies against IdU and pHH3 (markers for cells in the late G2-M phases) to label APs (apical surface-located IdU+/pHH3+) and BPs (basally located IdU+/pHH3+, filled arrows), which already passed through S phase and entered into late G-M phases. (B) Proportion of BPs, which entered the late G2-M phases in TSA-treated cortices is smaller compared with that in Veh-treated cortices. Values are presented as mean ± SEM (*P <0.05, **P <0.01, ***P <0.005). Abbreviations: VZ, ventricular zone; SVZ, subventricular zone; IZ, intermediate zone; CP, cortical plate. Scale bars = 50 μm.

**Fig. S7.**
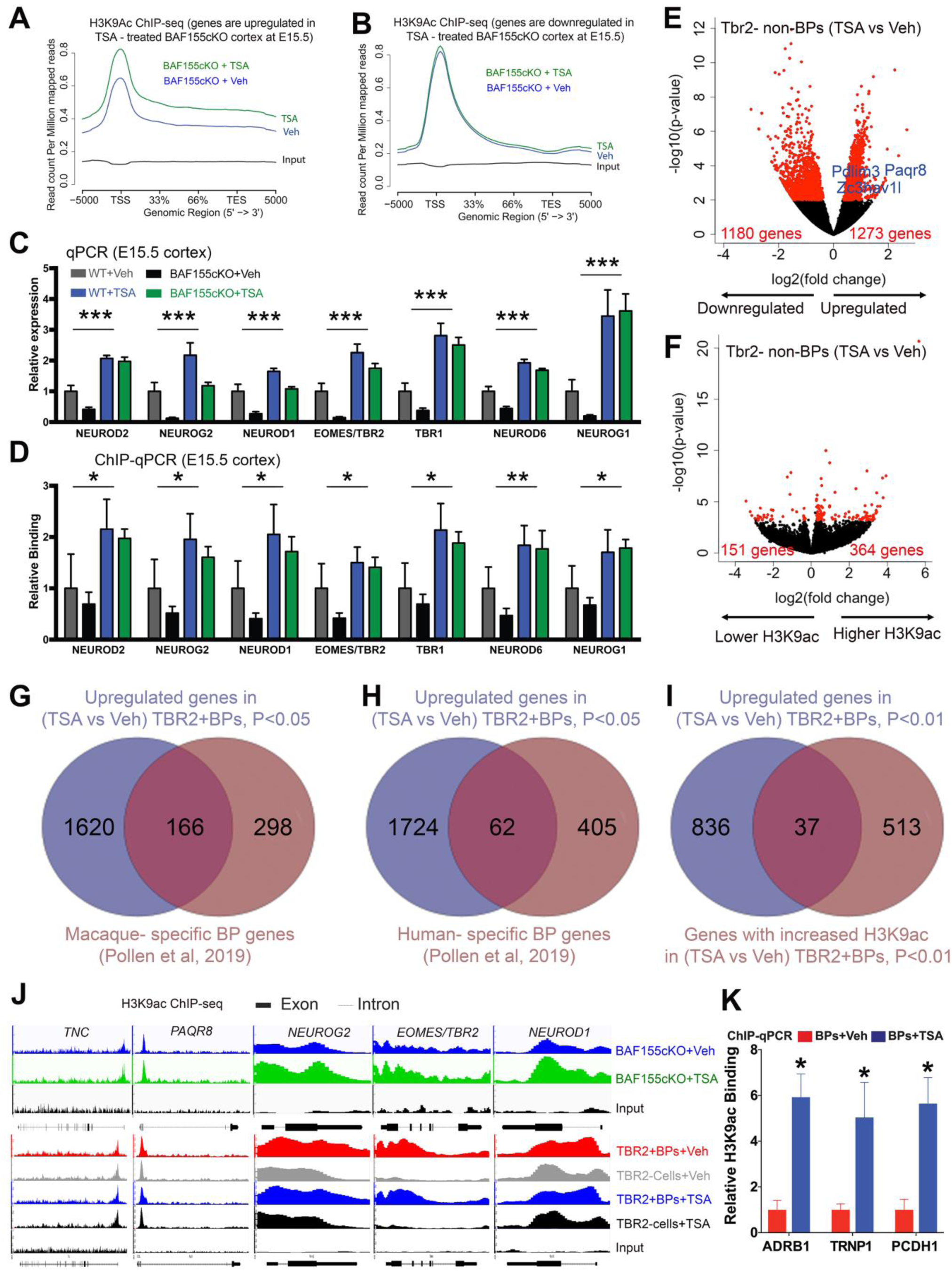
HDAC inhibition causes H3K9ac-linked upregulation of gene expression in BAF155cKO developing cortex, TBR2+ BPs and TBR2-cells. (A, B) H3K9ac is increased at loci of upregulated genes (A), but not at those of downregulated genes (B) in TSA treated BAF155cKO cortex. (C, D) Graphs of qPCR to examine gene expression (C) and ChIP-qPCR to quantify H3K9ac level (D). TSA treatment leads to increase in H3K9ac levels at promoters of neurogenesis-related genes (D) and their expression (C) both in WT and in BAF155cKO. ChIP-qPCR and qPCR results for a selection of genes are shown (WT_Veh: n = 3, WT_TSA: n = 4, BAF155 cKO_Veh: n = 4, BAF155 cKO_TSA: n = 4; general TSA effect: mixed linear model *p*-value < 0.001; general effect of BAF155 knockdown: Two-way ANOVA WT_Veh vs BAF155 cKO_Veh p < 0.01). (E, F) Volcano plots showing statistically significant changes (Paired Student’s *t*-test < 0.01, FC > 1.2) visualized by our RNA-Seq (E) and H3K9ac ChIP-Seq (F) analyses of TBR2-cells in TSA vs. Vehicle experiments (see also Fig. 5C/D for TBR2+ BPs). (G, H) Overlap between the upregulated genes in TSA-treated TBR2+ BP genes and BP/IP genes, which were recently identified specifically for macaque (G), and human (H) (Pollen et al., 2019). Notably, the TSA treatment provoked expression of a large set of macaque BP genes and in less extend, human BP genes in TBR2+ BPs in the developing mouse cortex. (I) Overlap between up-regulated genes in RNA-seq and genes with increased H3K9ac in ChIP-seq studies of TBR2+ BPs upon the TSA treatment. (J) Genome browser views of the distribution of H3K9ac along representative BP-enriched genes in BAF155cKO cortex, TBR2+ BPs, TBR2-cells, which were treated with either TSA and or vehicle as control. It should be noted that the bulk level of H3K9ac at their loci was unaltered in TBR2+ BPs and TBR2-cells in response to TSA treatment. In BAF155cKO cortex, the increased level of H3K9ac at these loci is due to expanded pool of BPs upon the treatment of TSA. (K) The increased level of H3K9ac at gene loci of ADRB1, TRNP1, PCDH1 in ChIP-seq upon TSA treatment were validated in ChIP-qPCR.

**Fig. S8.**
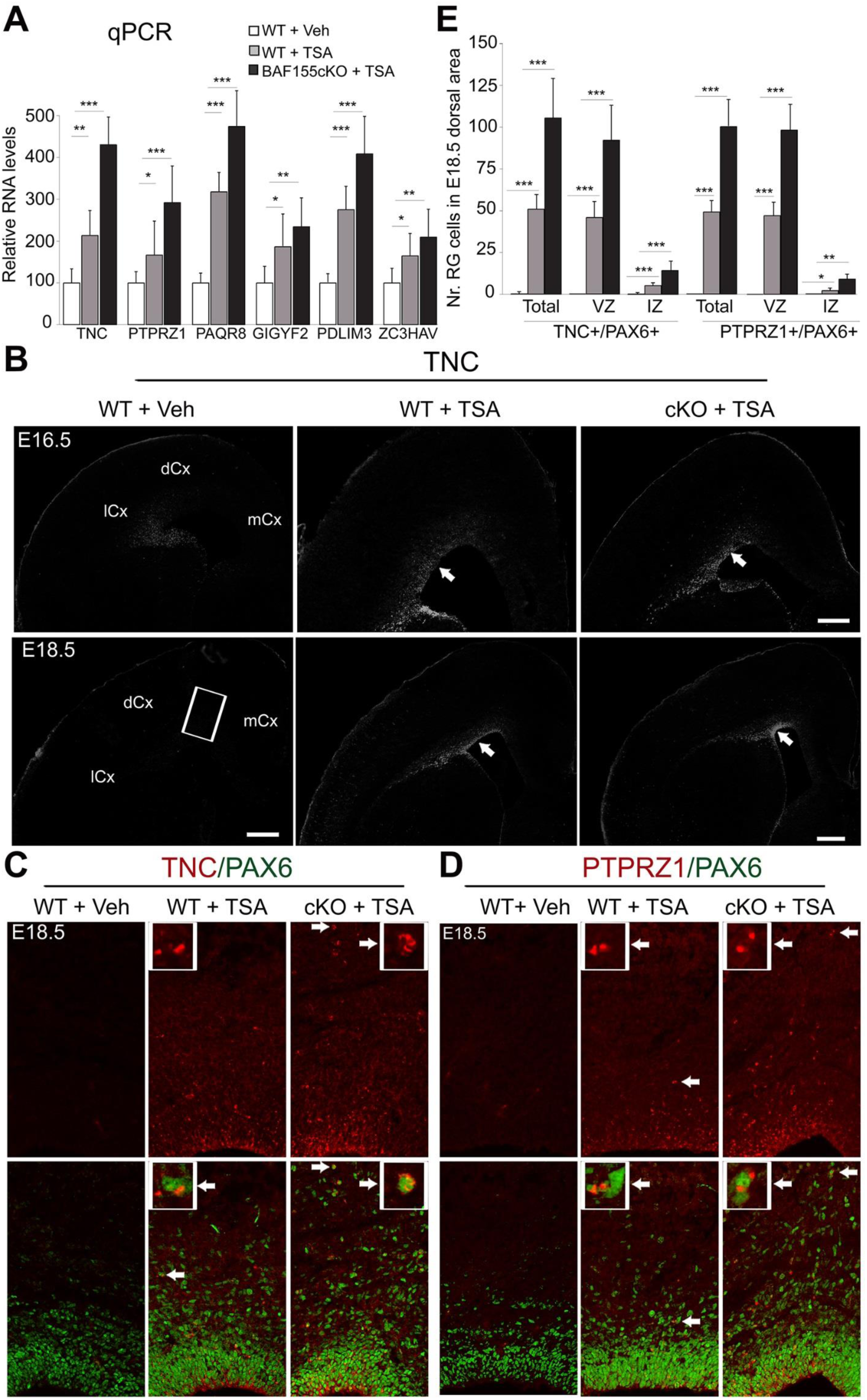
HDAC inhibition causes upregulation of bRG-enriched genes in developing mouse cortex. (A) qPCR analysis was used to confirm the upregulated expression of human-enriched bRG markers in TSA-treated cortex as compared to vehicle-treated control. (B-E) Among bRG-enriched genes, expression of TNC and PTPRZ1 have been well characterized in the developing cortex of both human and mouse (Pollen et al., 2015). TNC and PTPRZ1 antibodies labeled subsets of APs and BPs/bRGs in the entire human cortex, whereas in mouse their expression was detected only in a sub-population of PAX6+ APs in the lateral cortex (lCx, B) (Pollen et al., 2015). The increased expression of TNC (B/C) and PTPRZ1 (D) in TSA-treated WT and BAF155cKO cortex at E16.5 and E18.5 was revealed by immunolabeling (B-D) and quantification (E). Notably, the expression of these human bRG markers PTPRZ1 and TNC was found in PAX6+ APs in VZ of dorsal cortex (dCX, white filled arrows in B) and PAX6+ bRGs (white filled arrows in C, D) in TSA-treated cortex. Uppers images show the expression in arrow-pointed cells in lower pictures at higher magnification. Values are presented as mean ± SEM (*P <0.05, **P <0.01, ***P <0.005). Abbreviations: l/d/mCx, lateral/dorsal/medial cortex. Scale bars = 100 μm.

**Fig. S9.**
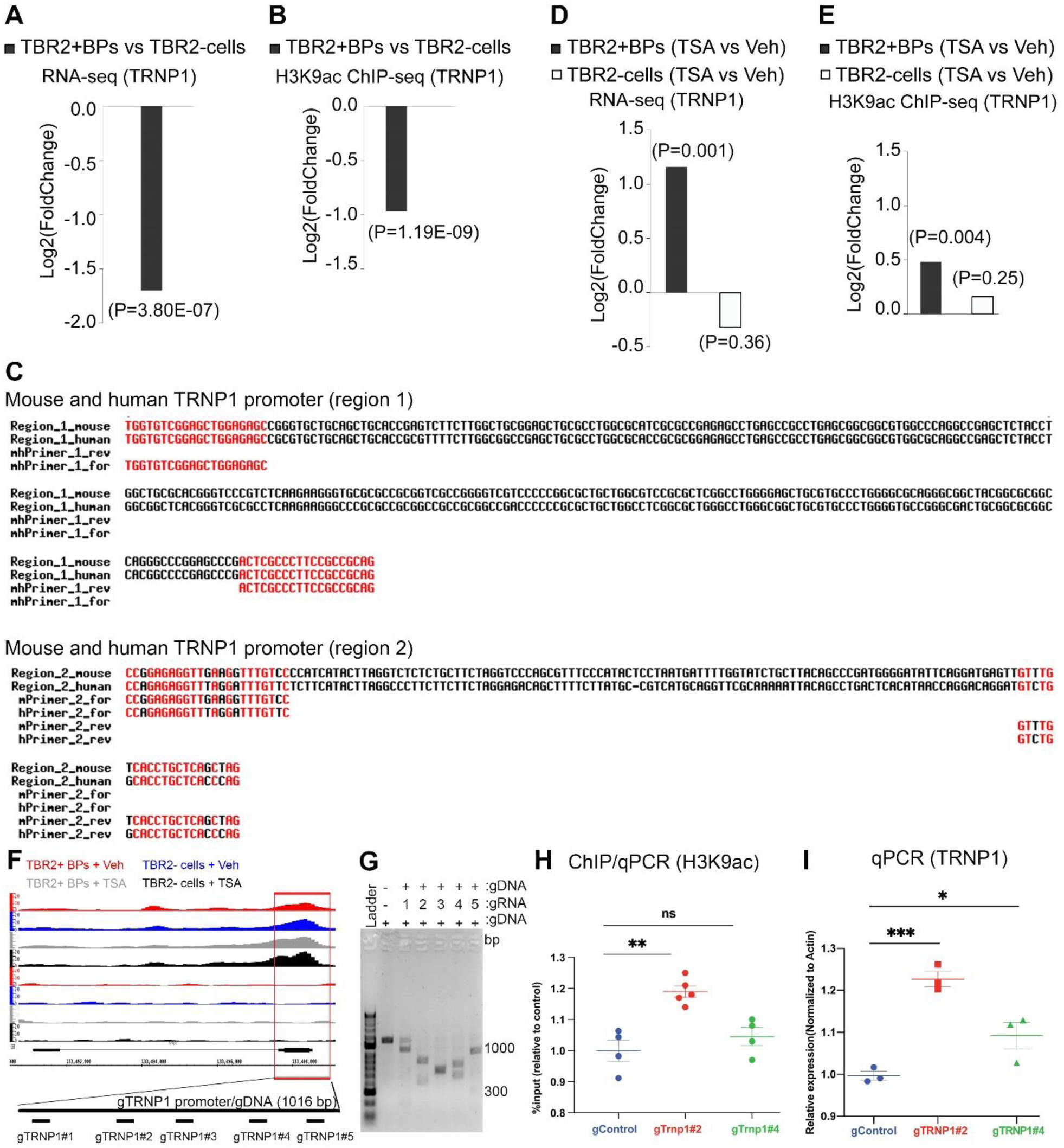
In vitro testing of TRNP1 gRNAs. (A/B) TBR2-cells have higher basal levels of H3K9ac at TRNP1 promoter (A) and higher basal expression (B) of this gene compared to TBR2+ BPs. (C) DNA sequence alignment for two ortholog regions of mouse (m) and human (h) TRNP1 promoter and primers, which were used in CHIP/qPCR experiment (see also Fig. 6C). Note that mhPrimer_1 set was used to amplify the region 1 of both mouse and human TRNP1 promoter. The mPrimer_2 and hPrimer_2 sets, which have similar sequence, were used to amplify the region 2. (D/E) TSA treatment significantly increases H3K9ac levels at TRNP1 promoter (D) and upregulates its expression (E) specifically in TBR2+ BPs but not in TBR2-cells. (F/G) Analysis of the Cas9 cutting efficiency guided by various gRNAs targeting the TRNP1 promoter. (F) Depiction of different gRNAs targeting various regions in TRNP1 promoter that were used for testing. (G) Agarose gel showing the cutting efficiency of each tested gRNA-Cas9 complex on a 1016-bp-long PCR product of TRNP1 promoter region. (H/I) ChIP/qPCR (H) and qPCR (I) analyses indicate that transfection of Neuro2A cells with gTRNP1#2 and #4 increases H3K9ac level at TRNP1 promoter (H) and upregulates its expression (I). Values are presented as means ± SEMs (*P <0.05, **P <0.01, ***P <0.005).

**Fig. S10.**
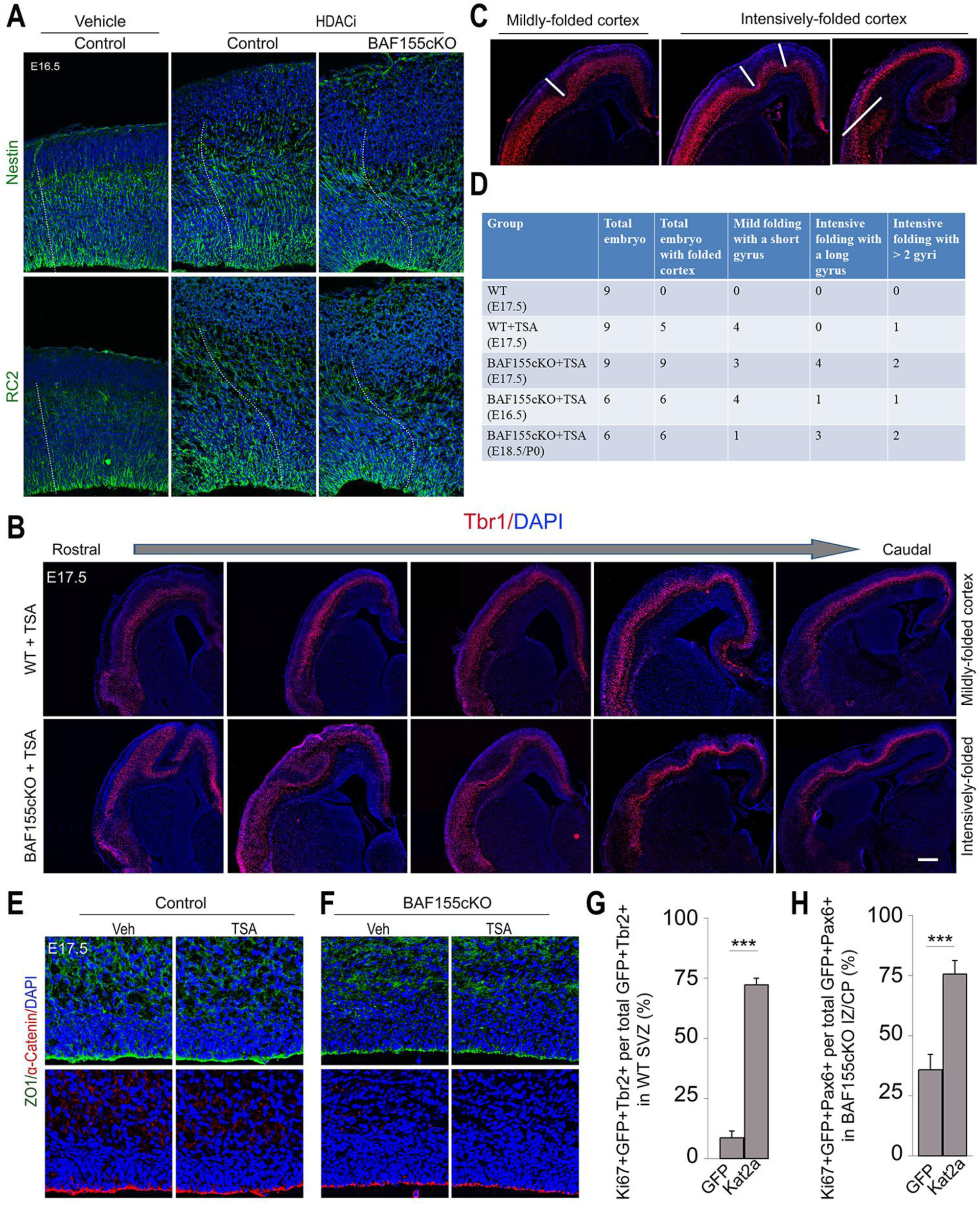
Augmented level of H3K9ac by HDAC inhibition and overexpression of KAT2A increase basal progenitor proliferation and can induce cortical folding. (A) Staining of RG fibers (NESTIN and RC2) in Veh-treated control, TSA-treated-control and TSA-treated BAF155cKO cortex revealed an increase and divergence of radial fibers at basal sides upon TSA treatment (radial processes indicated by white-dashed lines on confocal images). (B) Cortical tissue was processed for TBR1 immunostaining. Examples with different forms of cortical folding were shown from rostral cortex to caudal cortex at E17.5. The mildly-folded and intensively-folded cortices were usually found in TSA - treated WT embryos and TSA - treated BAF155cKO embryos, respectively. (C) The folding in length was measured from the pial surface to the end of the innervated point outlined by white lines. Cortical folding phenotypes were scored as a mild folding being shorter than 200µm in length (leftmost image in B) and intensive folding with a gyrus deeper than 200µm from the surface or with more than two gyri (right images). (D) Overview of cortical phenotypes observed after TSA or Veh-treatment. In total, nine Veh-treated control, nine TSA-treated control and nine TSA-treated *BAF155cKO* brains at E17.5, six brains at E16.5 and six brains at E18.5/P0 of TSA-treated BAF155cKO mutants were used to examine folding phenotypes. Six coronal sections per brain covering the rostral, middle-level and caudal cortex were processed for TBR1 IHC. Cortex was scored as “mild folding” or “intensive folding”, if at least two sections met the above criteria. (E/F) Staining of ZO1 and α-Catenin in treated control (E), and BAF155cKO cortex (F) revealed an intact VZ basement membrane upon TSA treatment. (G-H) E13.5 cortices of WT or BAF155cKO embryos were electroporated with eGFP or KAT2A-ires-eGFP, followed by quantification of TBR2+/KI67+ proliferating BPs/IPs (G) and PAX6+/KI67+ proliferating BPs/bRGs (H) at E15.5. Statistical analyses revealed that the overexpression of KAT2A led to increase the proliferation of BPs. Scale bars = 100 μm.

- **Table S1:** Differential gene expression between TSA- and vehicle-treated BAF155cKO embryonic cortex (as a Supplemental Spreadsheet).

- **Table S2:** Differential binding of H3K9ac at promoters between TSA- and vehicle-treated BAF155cKO embryonic cortex (as a Supplemental Spreadsheet).

- **Table S3:** Differential gene expression between TBR2+ BPs in TSA- and vehicle-treated embryonic cortex (as a Supplemental Spreadsheet).

- **Table S4:** Differential binding of H3K9ac at promoters between TBR2+ BPs in TSA- and vehicle-treated embryonic cortex (as a Supplemental Spreadsheet).

- **Table S5:** Differential gene expression between TBR2-cells in TSA- and vehicle-treated embryonic cortex (as a Supplemental Spreadsheet).

- **Table S6:** Differential binding of H3K9ac at promoters between TBR2-cells in TSA- and vehicle-treated embryonic cortex (as a Supplemental Spreadsheet).

- **Table S7:** Comparisons of the upregulated genes in TSA-treated TBR2+ BP genes and BP/IP genes, which were recently identified specifically for macaque and human (Pollen et al., 2019) (as a Supplemental Spreadsheet).

- **Table S8:** List of primers for qPCR and ChIP-qPCR (as a Supplemental Spreadsheet).

- **Table S9:** statistical analyses (as a Supplemental Spreadsheet).

